# In Silico Modeling of Virus Particle Propagation and Infectivity along the Respiratory Tract: A Case Study for SARS-COV-2

**DOI:** 10.1101/2020.08.20.259242

**Authors:** Dixon Vimalajeewa, Sasitharan Balasubramaniam, Donagh P. Berry, Gerald Barry

## Abstract

Respiratory viruses including Respiratory syncytial virus (RSV), influenza virus and cornaviruses such as Middle Eastern respiratory virus (MERS) and SARS-CoV-2 infect and cause serious and sometimes fatal disease in thousands of people annually. It is critical to understand virus propagation dynamics within the respiratory system because new insights will increase our understanding of virus pathogenesis and enable infection patterns to be more predictable *in vivo*, which will enhance targeting of vaccines and drug delivery. This study presents a computational model of virus propagation within the respiratory tract network. The model includes the generation network branch structure of the respiratory tract, biophysical and infectivity properties of the virus, as well as air flow models that aid the circulation of the virus particles. The model can also consider the impact of the immune response aim to inhibit virus replication and spread. The model was applied to the SARS-CoV-2 virus by integrating data on its life-cycle, as well as density of Angiotensin Converting Enzyme (ACE2) expressing cells along the respiratory tract network. Using physiological data associated with the respiratory rate and virus load that is inhaled, the model can improve our understanding of the concentration and spatiotemporal dynamics of virus.

## 1 Introduction

Respiratory viruses are among the most transmissable viruses in the world and create a global health burden that contributes to thousands of illnesses and deaths annually. According to the American Center for Disease Control (CDC), influenza has killed between 12,000 and 61,000 people annually and caused approximately 460,000 hospitalisations per year in the United States. Respiratory syncytial virus (RSV) is is estimated to cause close to 33 million cases of respiratory disease each year globally, with approximately 3 million hospitalisations [1]. Currently, the world is grappling with a major pandemic caused by a coronavirus, SARS-CoV-2, that also targets the lungs, has infected many millions of people, killed approximately 700,000 thus far and has caused unprecedented upheaval to the world, both socially and economically.

Once a respiratory virus enters the nose or mouth, it can move down the throat towards the lungs but much of the virus will deposit on the walls of the upper respiratory tract and infect cells along the way, leading to amplification of the initial virus load. The dispersion of virus particles throughout the respiratory system (i.e., the virus load along the respiratory tract) depends on the respiratory pattern (i.e., inlet airflow rate), the concentration of virus particles in the inhaled air, also known as exposure level, and the distance that they can travel through the respiratory system. In addition, the bifurcation geometry of the airway as well as the distribution of cell surface receptors that the virus can bind to also change the virus dispersion along the respiratory tract. For example, in the case of SARS-CoV-2, the virus is known to use the Angiotensin Converting Enzyme (ACE2) receptor to bind to and enter cells [2]. Therefore, the density of ACE2-expressing cells in the respiratory tract is a critical factor in the distribution of virus. Since cells expressing ACE2 are not spread evenly throughout the respiratory system, the areas that are expressing ACE2 are more vulnerable to SARS-CoV-2 infection, thus creating a heterogeneous concentration distribution across the respiratory system. The virus concentration at any point in the respiratory system, therefore, will depend on the replication location as well as replication rate, which can result in either further propagation into the airway or potential transmission to other individuals through exhalation [3].

The evolution of the infection and the spreading pattern along the respiratory system can be characterized and predicted effectively if the viral concentration dynamics (i.e., virus propagation, deposition, infection and reproduction) can be characterized using modeling and simulations along with critical parameters that influence the system. This knowledge can provide new tools to improve personalized patient care, and support biotechnological and pharmaceutical research to design effective therapeutic drugs. From a personalization perspective, by being able to predict how the virus will impact a patient based on known criteria, medical staff will be more able to make timely, informed and accurate decisions to prevent development of severe symptoms that can worsen the patient’s condition. In the Pharmaceutical industry, this model can benefit the development of drugs by, for example, allowing the *in silico* screening and characterization of inhibitory drugs prior to costly *in vivo* trials.

The main aim of this study was to develop a generic computational model for characterizing the dynamics and the evolution of infectious respiratory diseases along the respiratory tract. First, the model integrates biophysical models of virus propagation and deposition within an airway flow system as well as infection properties such as the host cell distribution and virus replication process. Second, we formulate the virus concentration levels along the branches of the respiratory tract. Moreover, the model also accounts for virus dispersion along the respiratory tract that can be impacted by factors such as the respiratory rate, viral exposure levels (i.e., the quantity of virus that is inhaled) and the virus particle size. Integrating these together, the computational model explores the changes in the virus propagation as it encounters advection-diffusion based air flow, before turning into pure diffusion towards the end of the lungs (e.g., alveoli regions). In the virus infection process, the study also considers the impact of the immune system and how it impacts on the overall concentration of the virus within the respiratory tract. The study uses SARS-CoV-2 as a case study to present how the derived model could be used to characterize the virus propagation dynamics in humans. The main contributions of this study are as follows:

1. Development of a temporal viral concentration model that can be used to analyze changes in virus dynamics as virus particles propagate along the branches of the airway in the respiratory system. This is achieved by accounting for the generation branching geometry of the respiratory tract, and integrating models of the air flow that affects to viral deposition and replication, and its impact on the virus spread distribution along the airway generations.
2. Model the variability in spatial and temporal dynamics of the virus particle evolution within the respiratory tract, by considering personalized patient parameters such as the breathing rate and dynamic viral pleomorphic size change, with respect to the variability in inhaled dosage.
3. Sensitivity analysis of the viral propagation along the generation branches considering factors such as the exposure level as well as the immune response.

## 2 Methods

The main steps in the process of deriving the computational model for the virus propagation dynamics along the respiratory tract are illustrated in Fig. 1. There are three main elements to the model include the concentration dynamics based on particle deposition, virus propagation under advection-diffusion air flow over the respiratory airways and the impact from the infection process (complete derivation of each of these processes are provided in the Supplementary Information (SI) 4.4).

**Figure 1:**
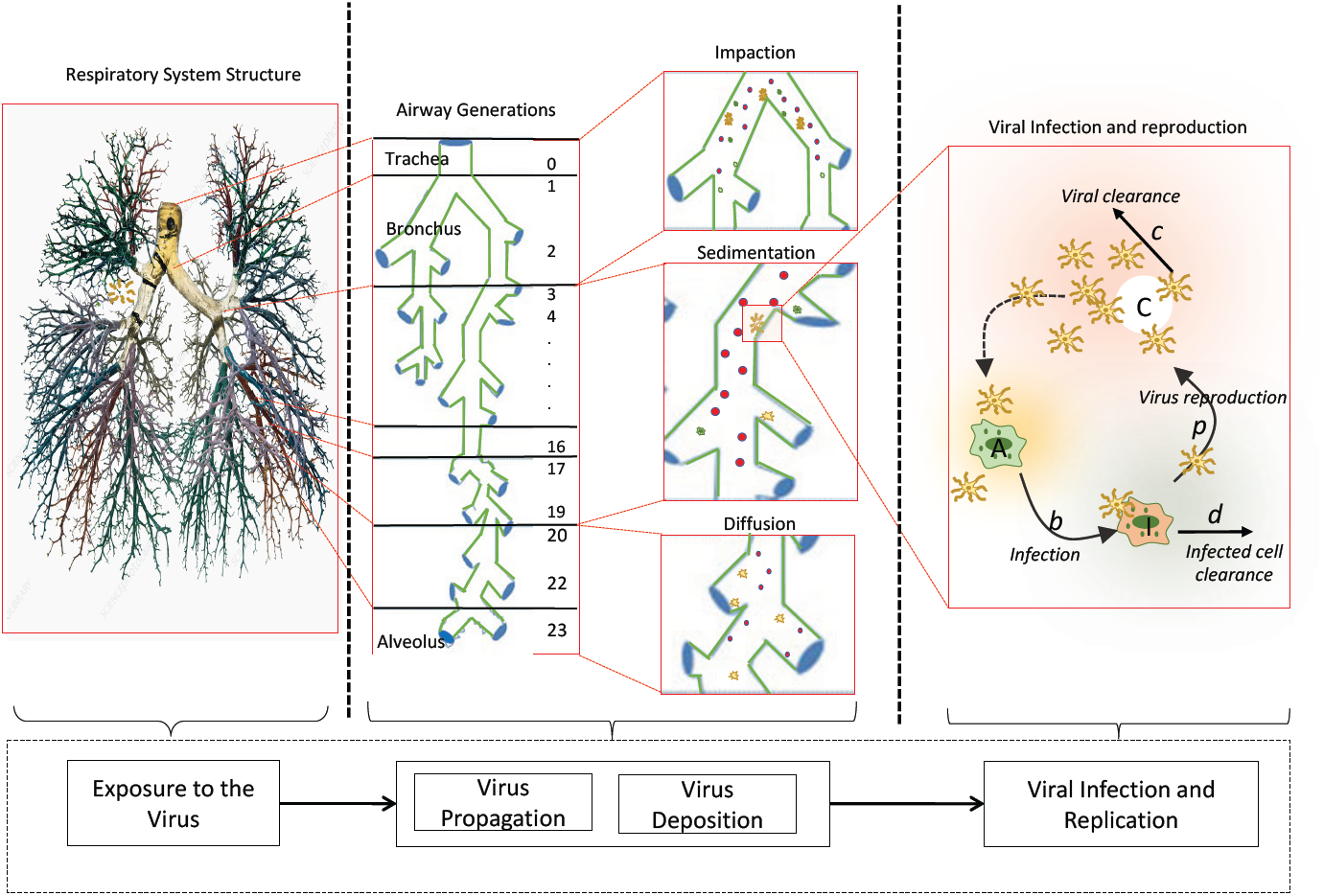
Computational model for exploring the evolution of viral propagation dynamics down the branch generation of the respiratory tract. The computational model considers the virus deposition and distribution of host cell density along the airway and their role in proliferating the virus to characterize the variability in viral concentration dynamics along the respiratory tract (the airway network figure used in this diagram is adapted from science photo)

### 2.1 Respiratory Tract Generation Model and Virus Concentration Dynamics

Fig. 2 depicts the respiratory airway generation structure that is used in our study to characterize the viral particles concentration and its distribution along the respiratory tract. The respiratory generation model in our study is taken from the study in [4] and includes the real physiological parameters, such as the diameter and length of each airway generation, and their values are provided in Table 1. The concentration of the viral particle in the *i*^*th*^ airway generation, 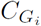, will depend on the quantity of virus that is in the previous airway generation (i.e., 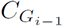). This means that the virus infection level (or the virus concentration) of the *G*_*i*_ airway generation depends on the virus infection level of the airway generation *G*_*i*−1_ and its propagation along the branch. Moreover, the viral particle deposition rate and the virus reproduction speed in the host cells along with the respiratory system geometry such as the airway length and diameter also contributes to the virus infection level.

**Table 1:** Physiological parameters of the respiratory airway generations taken from [4].

| Generation number | $G_0$ | $G_1$ | $G_2$ | $G_3$ | $G_4$ | $G_5$ | $G_6$ | $G_7$ | $G_8$ | $G_9$ | $G_{10}$ | $G_{11}$ |
| --- | --- | --- | --- | --- | --- | --- | --- | --- | --- | --- | --- | --- |
| | $G_{12}$ | $G_{13}$ | $G_{14}$ | $G_{15}$ | $G_{16}$ | $G_{17}$ | $G_{18}$ | $G_{19}$ | $G_{20}$ | $G_{21}$ | $G_{22}$ | $G_{23}$ |
| Length ($L$ cm) | 12 | 4.76 | 1.9 | .76 | 1.27 | 1.07 | .9 | .76 | .64 | .54 | .46 | .39 |
|  | .33 | .27 | .23 | .2 | .165 | .14 | .12 | .099 | .083 | .07 | .059 | .05 |
| Diameter ($d_L$ mm) | 1.8 | 1.22 | .83 | .56 | .45 | .35 | .28 | .23 | .186 | .154 | .130 | .109 |
|  | .095 | .082 | .074 | .061 | .06 | .054 | .05 | .047 | .045 | .043 | .041 | .04 |

**Figure 2:**
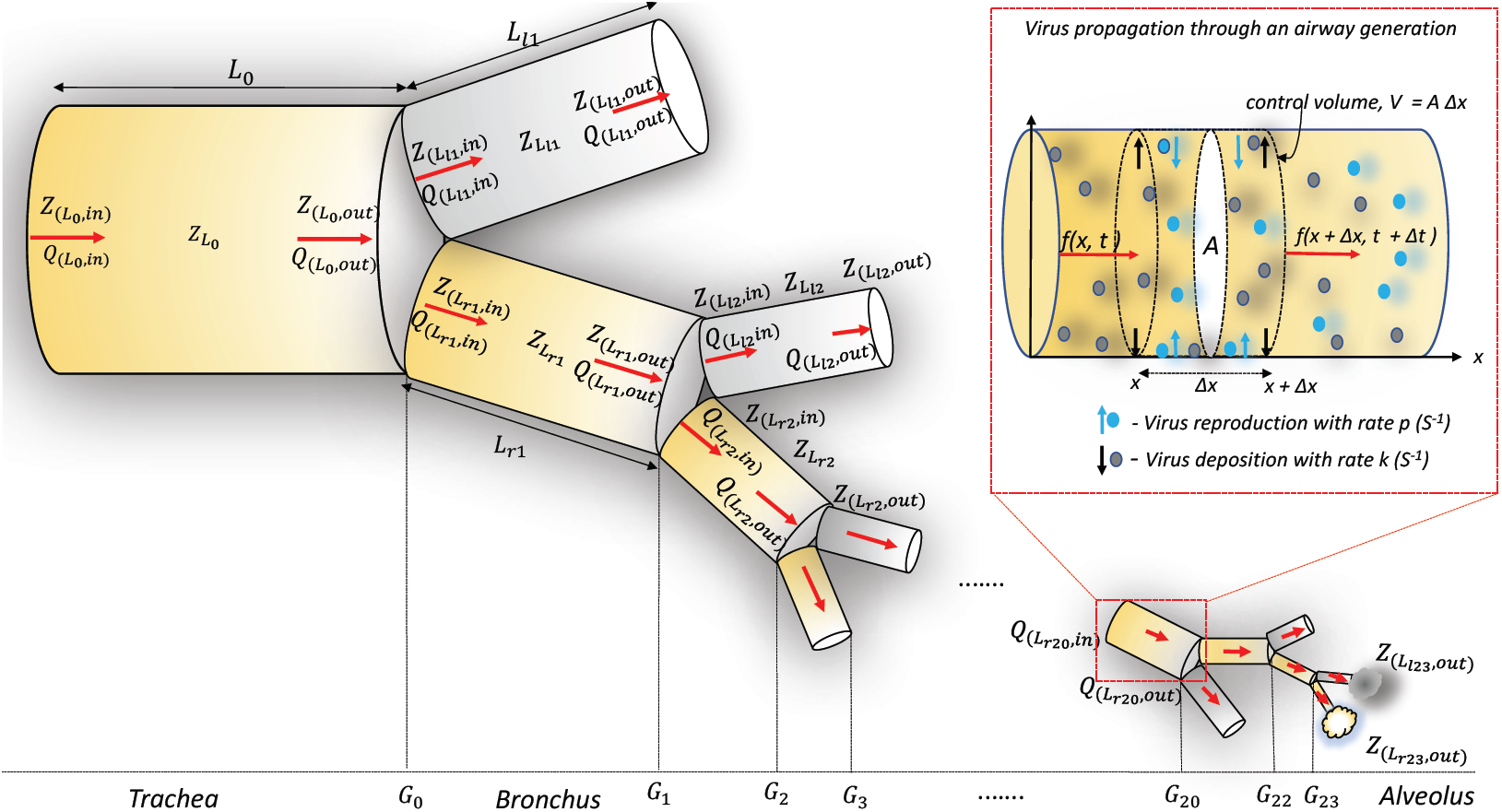
Bifurcation of the respiratory (left - *l* branch and right-*r* branch) airways; the subscript notation *in* and *out* are used to denote the inlet and outlet pressure (*P*), airflow rate (*Q*) and impedance (*Z*) in each airway and virus particle propagation through a control volume of an airway with deposition rate *k* and reproduction rate *p*.

By taking these factors into account, this section presents how the advection-diffusion mechanism of the virus particle propagation can be utilized to compute the concentration over an airway generation *G*_*i*_. When the virus particles travel down the respiratory tract under the advection-diffusion mechanisms, they deposit on the airway walls and bind onto ACE2-expressing cells to initiate infection and subsequently the replication process. Considering the viral infection and replication process depicted in Fig. 2 and the mass balance of the virus particles in the control volume, the change in the virus concentration *G*_*i*_, where *i* represents an airway generation, is represented as

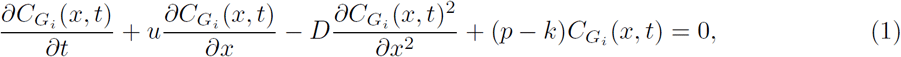

where the generation number *i* = 0, 1, 2,…, 23, *k* and *p* are the virus deposition and reproduction rates, respectively, *D* is the air diffusion coefficient and *x* stands for the direction of the virus propagation (i.e., downward of the respiratory tract). Given the initial inlet virus concentration that a person consumes through the mouth or nose is 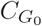 (i.e., 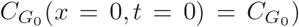), the solution of Eq. 1 can be derived as

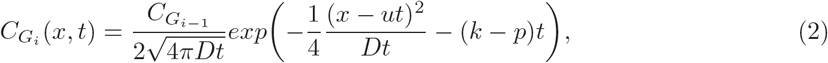

where 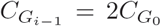 only at *t* = 0 as 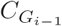 divides equally into two when the virus enters the branches of the *G*_*i*_ generation for *i* = 1, 2, 3, *…*, 23.

At steady state, 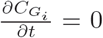 and *u* is also a constant. This means that Eqn. 1 can be simplified into *Da*^2^ − *ua* − (*k* − *p*) = 0 with 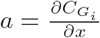. This is a quadratic expression and thus, the solution is 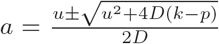. Since concentration 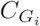 declines over time and *C ≥* 0, 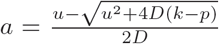. Using this value of *a*, the virus concentration at the steady state is expressed in Eq. 3 (see SI for more details 4.4). The change in the virus concentration along an airway that includes the effects of virus reproduction and their deposition on the airway walls is represented as

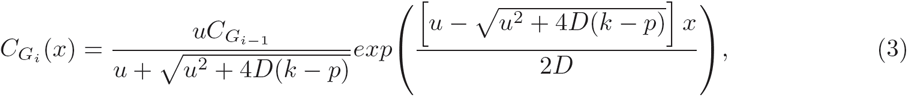

for *i* = 0, 1, 2, *…*, 23.

Next, the following three subsections describe the methods for computing the air flow velocity (*u*), virus deposition rate (*k*) and virus reproduction rate (*p*), which are required for determining the virus concentration using Eq. 3.

### 2.2 Virus Propagation over the Airways

In order to calculate air flow velocity *u* required to determine the virus concentration in Eq. 3 along the airway generation branches, we take into account the bifurcation transformation between the large airways that separates into smaller branches with reduced lengths and diameters as illustrated in Fig. 1. Suppose an airway of length *L*_0_ branches into left and right branches of length *L*_*l*_1 and

*L*_*r*_1 as illustrated in Fig. 2. Based on the conservation of air flow, at the first bifurcation (i.e., *G*_0_), the conditions 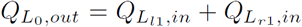 and 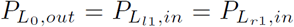 holds at the bifurcation junction; where *Q* is the flow rate, *P* is the pressure, *L*_*l*1_ and *L*_*r*1_ are respectively the lengths of the left (*l*) and right (*r*) branches, and the numbers *l* and *r* refers to the airway generation number. In addition, the subscript *in* and *out* stands for the inlet and outlet of the branch. When air flows down the airway, there is a drop in pressure as well as resistance against the air flow. We adapt these properties and model the flow using transmission line circuit theory, where its equivalence is voltage drop in response to changes in the current flow [5]. Hence, following the concepts from transmission line theory, the impedance (*Z*) at the outlet of the parent airway 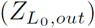 is related to the impedance at the inlet of the branching airways as 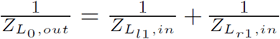. This results in the impedance at the inlet being expressed as 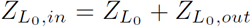, where 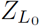 is the impedance of the airway (4.4 for the derivation of 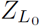). Based on this, the inlet air flow rates to the branches and the outlet pressure of the parent airway can be represented as

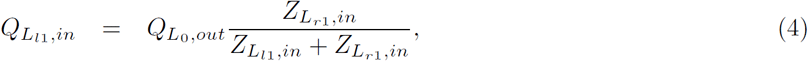

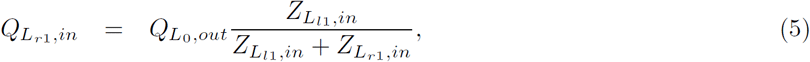

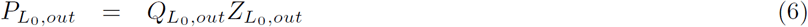

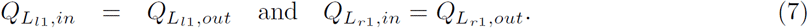

To compute the airflow rates, as well as the velocity profiles over the respiratory tract, we use a backward calculation procedures, which starts with the impedance at the outlet of the last airway (i.e., 23rd generation’s 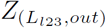 and 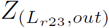 as represented in Fig. 2). As an example, the procedure for the 23rd and 22nd generation airflow rate is as follows:

1. The impedance of the 23rd airway generation, *Z*_23_, as well as the impedance at the inlet of this airway, *Z*_(23,*in*)_, is computed (see SI for formulas 4.4).
2. The impedance at the outlet of the 22nd airway generation, *Z*_(22,*out*)_ is then computed based on the impedance values of step 1 (23rd generation).
3. Step 1 is repeated to compute the impedance of the 22nd airway generation, *Z*_22_, as well as the impedance at the inlet, *Z*_(22,*in*)_.

These steps are repeated up to the initial airway generation (i.e., 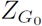 as represented in Fig. 2) to compute the impedance of each airway generations. Then, the flow rate *Q* into each airway generation is computed using Eqns. 4 and 5. Having derived the flow rate into the airway, the air flow velocity *u* is then computed using the expression *Q* = *Au*, where *A* = *π*(*d*_*L*_*/*2)^2^ is the cross sectional area of the airway of diameter *d*_*L*_. The real physiological parameters, length (*L*) and diameter of the airway generations (*d*_*L*_) given in Table 1 (taken from [4]) are used for computing the air flow velocity of each airway generation.

The air enters into the respiratory tract under external forces that includes both turbulence as well as gravity, and the impact of these forces gradually decreases once we go deeper into the lung. Hence, the propagation of particles contained in the airflow can be determined using the advection-diffusion property [6]. The advection mechanism is the dominant force that facilitates the propagation of particles and their deposition along the upper airways due to the high airflow velocity, which in turn, causes turbulence in the airflow. This differs from the lower branches within the respiratory track, which is dominated largely by pure diffusion.

#### 2.2.1 Particle deposition rate

The virus deposition (*k*) mainly takes place under three main mechanisms, impaction (*k*_*I*_), sedimentation (*k*_*S*_) and diffusion (*k*_*D*_). Hence, the final particle deposition rate *k* is computed as the sum of all deposition rates and is represented as

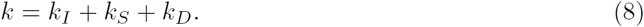

Details for *k*_*I*_, *k*_*S*_ and *k*_*D*_ are as follows ([4], [7], [8]):

1. **Impaction (***k*_*I*_ ): Virus deposition due to impaction takes place in response to virus particles which cannot flow evenly along their trajectory due to their inertia and surroundings. These barriers increases with particle size and flow rate and mostly occurs in the upper airways due to high velocity. In our study, we use the following formula to compute *k*_*I*_ [4].

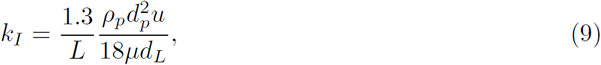

where *L* is the airway length, *ρ*_*p*_ is the virus particle density, *d*_*p*_ is the virus particle diameter, *µ* is the air viscosity and *d*_*L*_ is the airway diameter.
2. **Sedimentation (***k*_*S*_): Sedimentation occurs when the virus particles move under the gravitational force, and they settle on the lower surface of the airways in the upper respiratory region, and is computed as [4]

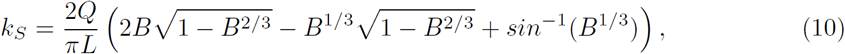

where 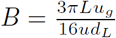 and *u*_*g*_ is the settling velocity of the virus particles. Settling velocity is known as the maximum velocity that a virus particle can achieve (i.e., velocity of a particle when the total external forces acting on it becomes zero). Fig. 3 depicts a virus particle of mass *m*, diameter *d*_*p*_ and density *ρ*_*p*_ experiencing an airflow of density *ρ*_*f*_ and velocity *u*_*f*_ at *t* = 0 and then propagates down with velocity *u*_*p*_ under three forces, which includes external gravity, drag and buoyancy. When the virus particle achieves its settling velocity *u*_*g*_ at *t* = *T* after traveling *L*_*T*_ distance, the particle moves at an arbitrary time *t*, where its relative velocity (velocity with respect to the airflow) can be expressed as *u*_*p,t*_ = *u*_*f*_ − *u*_*p*_. The forces (external force *F*_*e*_, drag force *F*_*D*_, and buoyancy force *F*_*b*_) acting on the particle, shown in Fig. 3, Eqn. 11 is derived by applying Newton’s second law, which is used to determine the settling velocity *u*_*g*_, and is represented as follows:

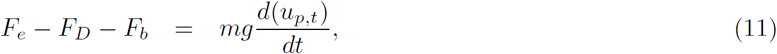

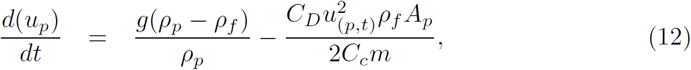

where *C*_*D*_ is the drag coefficient, *C*_*c*_ is the slipping coefficient, *g* is the acceleration of gravity and *A*_*p*_ is the cross sectional area of the virus particle (see SI for more information about deriving *u*_*g*_ 4.4).
3. **Diffusion (***k*_*D*_): Diffusion-based particle deposition acts on the virus particle resulting from Brownian motion. This increases in response to decreasing particle size *d*_*p*_ and and flow rate *Q*_0_ [4], and is represented as

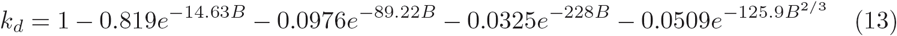

where 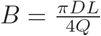, *D* is the diffusion coefficient, *Q* is the flow rate, and *L* is the airway length.

**Figure 3:**
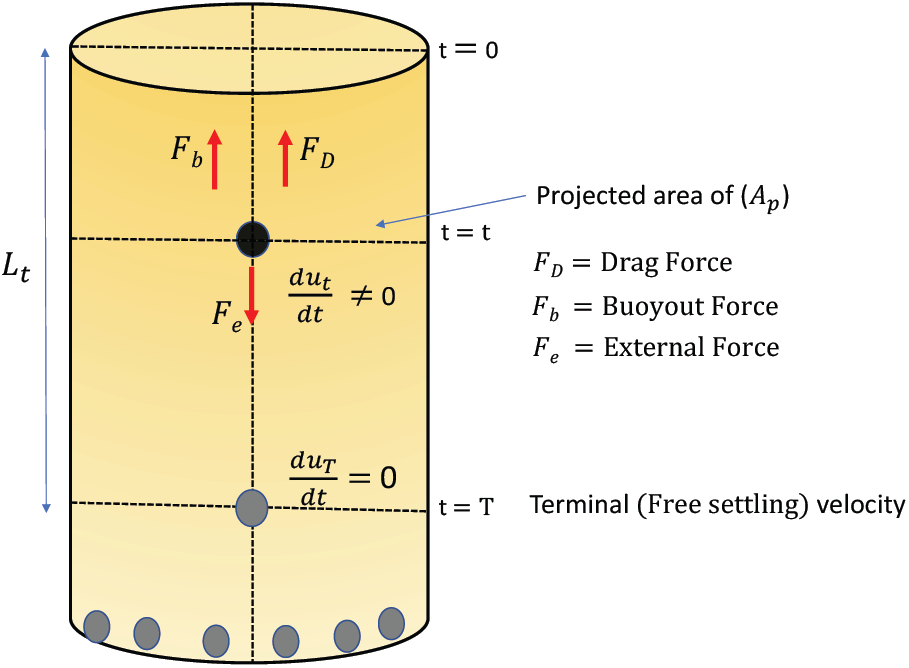
Virus particle propagation under gravity, achieving settling velocity and then deposition on the airway walls.

### 2.3 Virus infection process

In the event of a virus infection, SARS-Cov-2 uses ACE2 as a cellular receptor, enabling entry into a number of different cells found in the respiratory tracts (e.g., tracheal and bronchial epithelial cells, type 2 pneumocytes, macrophages) [9]. The infection also stimulates increase in ACE2 expression and this in turn accelerates the virus infection process. In addition, the viral infection triggers the transcription of interferon stimulated genes which impact on the viral reproduction as well as as promoting an inflammatory response [10] and this is followed by cellular apoptosis in response to the infection. Based on previous studies related to the infection of diseases such as HIV and influenza [11], the basic model used to describe this viral infection process is given in Eqn.14-16. This model expresses the inter-relationship between the ACE2-expressed cells (*A*) (also generally known as susceptible cells), infected cells (*I*) and virus (*C*), and is represented as

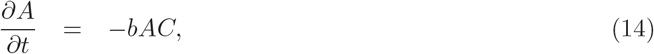

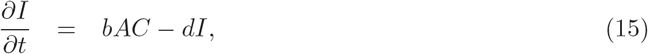

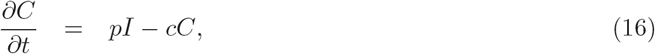

where *b* ((Copies/ml)^−1^ day^−1^) and *d* (day^−1^) are respectively the virus infection with susceptible cells and infected cell clearing (due to immune response) rates and *p* ((Copies/ml)^−1^ day^−1^ cell^−1^) and *c* (day^−1^) are the virus production rate and clearing rates, respectively. In this study, the solution of this system of equations expresses the change in *A, C* and *I* concentrations over time and is derived numerically by using Ordinary Differential Equation Solvers(ODE) [12].

Finally, the airflow velocity *u*, the virus deposition rate *k* and the virus replication rate *p* computed respectively using Eqns. 4-7, Eqn. 8 and Eqns. 14-16 are used to determine the virus concentration in each airway generation 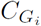 (*i* = 0, 1, *…*, 23) by using Eqn. 3.

## 3 Results

This section mainly presents the spatial and temporal viral dynamics over the respiratory tract. The computational outcomes presented below are generated by using the parameter values listed in Table 2, unless otherwise mentioned for specific analysis. In our results analysis, the impact of the airflow rate and flow velocity on the deposition of viral particles on the airway walls is explored. The virus infection process based on the deposition process is studied and this is followed by the propagation dynamics of the viral particles along the respiratory tract. Finally, this analysis explores the impact of several parameters such as the respiratory rate, immune response and exposure level on the dynamic propagation of virus particles through the respiratory tract.

**Table 2:** Parameters and notations along with their values used in simulations

| Parameter | Value |
| --- | --- |
| **Airflow characteristics** |  |
| Air density ($\rho_f$) | $1.2 \times 10^{-4} (gcm^{-3})$ |
| Air viscosity ($\mu$) | $1.81 \times 10^{-4} (gcm^{-1}s^{-1})$ |
| Inlet airflow ($Q_0$) [4] | $30 (lmin^{-1})$ |
| Acceleration of gravity ($g$) | $980 (cms^{-2})$ |
| Mean free path of gas molecules ($\lambda$) [13] | $0.066 \times 10^{-4}$ |
| Boltzmann constant ($K_B$) | $1.38 \times 10^{-16}$ |
| Room Temperature ($T$) | $298 (K)$ |
| Airway Length ($L$) and diameter ($d_L$) [4] | Table 1 |
| **Virus characteristics** |  |
| Inlet virus concentration ($C_0$) [2] | $1 \times 10^7 (Copies/ml)$ |
| ACE2 concentration ($A$) [9] | $\mathcal{N}(5.83, .71) (Copies/ml)$ |
| Diameter of virus particle ($d_p$) [14] | $60-140 (nm)$ |
| Density of virus particle ($\rho_p$) [15, 16] | $1.18 (gcm^{-3})$ |
| Virus reproduction rate ($p$) [2] | $8.2 (day^{-1})$ |
| Virus clearing rate ($c$) [2] | $0.6 (day^{-1})$ |
| Virus infection rate with susceptible cells ($b$) [2] | $3.9 \times 10^{-7} (Copies/ml)^{-1} day^{-1}$ |
| Infected cell clearing rate ($d$) [2] | $4.71 (day^{-1})$ |

### 3.1 Virus propagation through the respiratory tract

In exploring the virus propagation along the respiratory tract, one of the most critical insights required is the change in airflow along the airway generations starting from the trachea to the alveolar regions. Here, we used the derived model in Eqn. 4 to compute the airflow rate and the velocity in each airway generation by using real physiological parameters such as the length and diameter of the airway generations [4]. Fig. 4a shows both the change in the airflow rate and velocity over the 23 generations when air enters through the nose or mouth at a rate of 30 *lmin*^−1^. The airflow rate reduces to nearly half its value as it progresses into higher airway generations due to the bifurcation of larger airways that are separated into smaller branches. The velocity shown in Fig. 4a also reduces with higher airway generations. This validates our belief stated in the previous section, that the dominant force that propagates the virus into the lungs is governed by the advection and diffusion process, and in particular in the upper and lower parts of the respiratory system. Fig. 4b depicts the virus deposition under sedimentation and impaction, and shows again that this is higher in the upper airways, while the particle deposition due to diffusion increases with increasing airway generation number.

**Figure 4:**
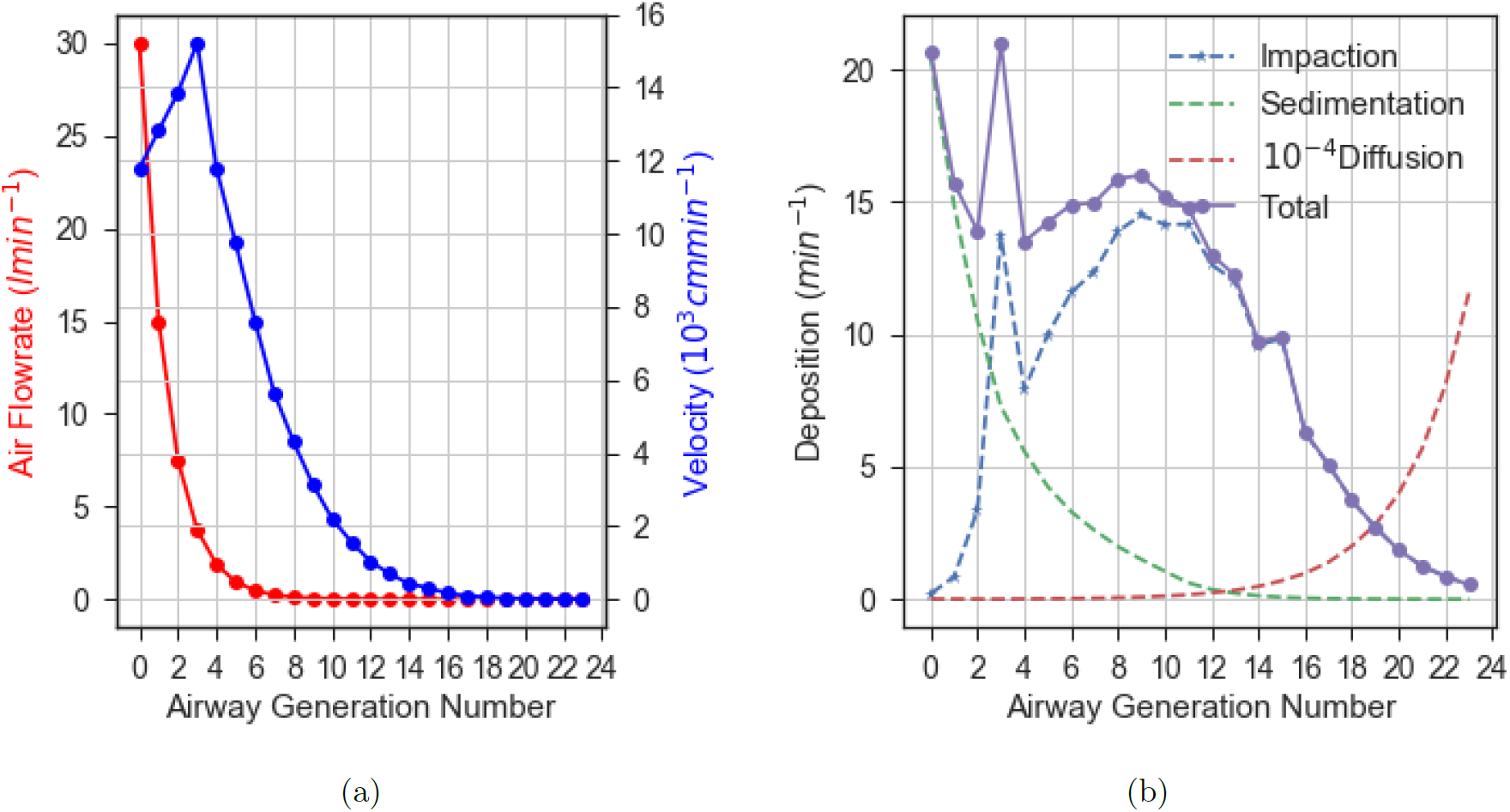
Change in airflow rate *Q* and velocity *u* along with deposition *k* over the airway generations in the respiratory system when the inlet airflow rate *Q*_0_ = 30*lmin*^−1^.

The virus shape is pleomorphic (i.e., changes shape) and is predominantly elliptical or spherical. Therefore, we also analyzed the impact of the virus particle shape and its propagation through the respiratory system based on changes in its diameter between 60–140 *nm* [14]. *In terms of velocity and deposition*, Fig. 5a(*I* − *IV*) and Fig. 5b(*I* − *IV*) depicts the impact of variability in the inlet airflow rate (*Q*_0_ = 15 − 60*lmin*^−1^) on the airflow velocity and virus deposition through the respiratory system. The results show that the velocity of the virus particles as well as their deposition rate *k* increases over the airway generations with increasing inlet airflow rate (i.e., increasing breathing rate). It can be observed further that the variability in the virus deposition rate over the first 15 airway generations is considerably higher compared to the airway generations in the alveolar region.

**Figure 5:**
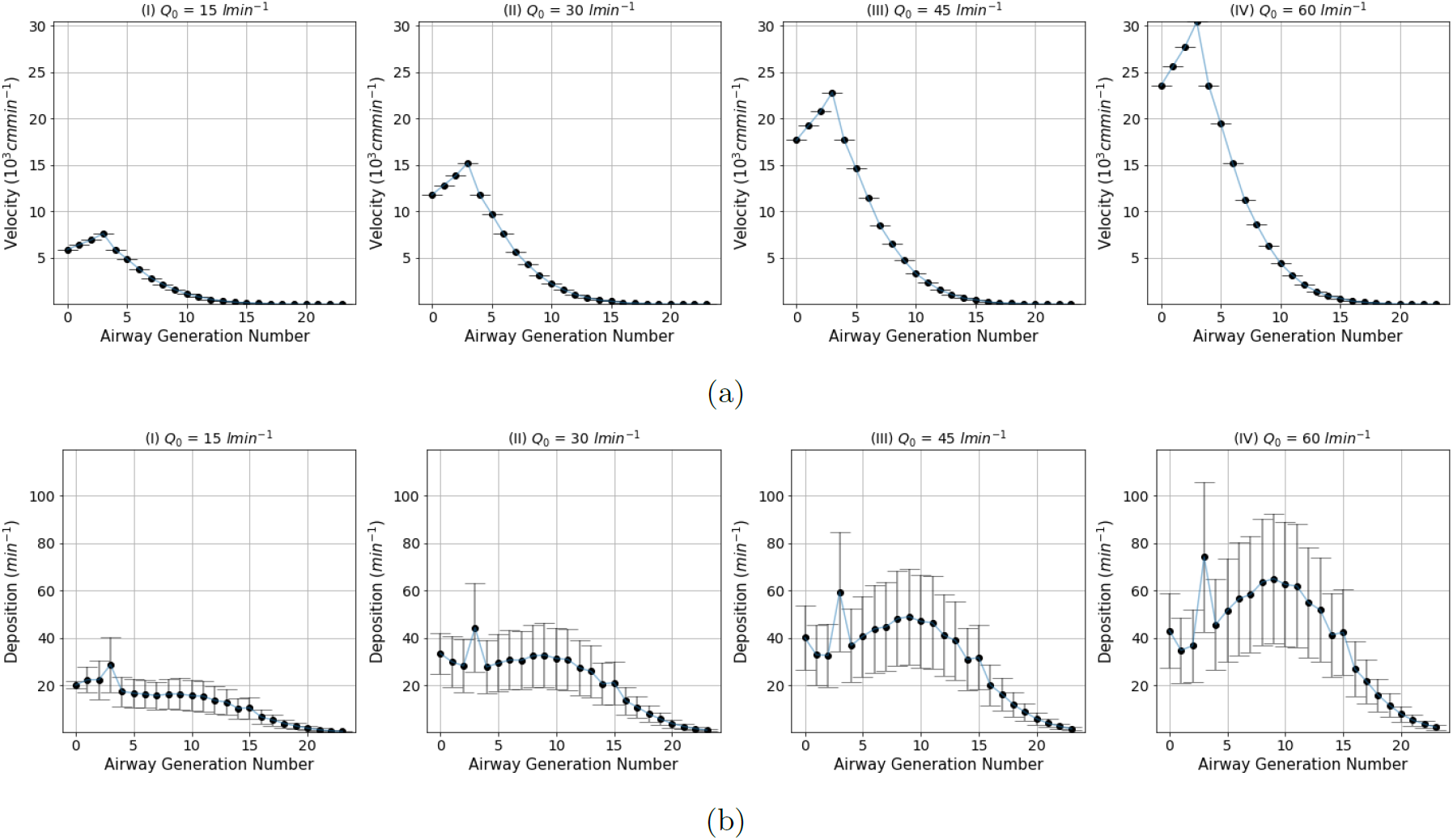
The impact of airflow rate *Q* and particle diameter *d*_*p*_ on the airflow velocity *V* and particle deposition rate *k*; **(a)**airflow velocity *V* and **(b)** deposition rate *k* with respect to variations in the airflow rate *Q* (15-60 *lmin*^−1^) and diameter of the virus particle *d*_*p*_ (70-130 *µm*), respectively.

In addition, the impact of changes in virus diameter *d*_*p*_(= 70 − 130*nm*) on the airflow velocity and virus deposition through the respiratory tract are also presented in Fig.4 in the SI 4.4.

### 3.2 Virus Concentration Dynamics Along the Airways

To understand the virus concentration dynamics as it flows along the airways, our analysis assumes a virus concentration of 10^7^*Copies/ml* (or Infectious particles/*ml*) entering the respiratory tract (i.e., *C*_0_) at a breathing airflow rate *Q*_0_ = 30*lmin*^−1^. In addition, we also assume the alveolar type-II and epithelial cell density that express ACE2 is higher around the mouth, nose and alveolar [6]. The higher quantity is achieved by deriving ACE2 concentration for each airways generation from the Gaussian distribution, where we averaged higher ACE2 expression (∼6 −7 *Copies/ml*) around the trachea and alveolar (i.e., ∼1 −2 and ∼21 −23 airway generations) compared to the other airway generations (∼4 −5*Copies/ml*), but with the same variance.

Fig. 6(*I*) shows the variability in the virus concentration computed using Eqn. 3, while Fig. 6(*II*) depicts the change in ACE2 expression levels over the airway generations. It can be observed that the abundance of virus is higher in the upper airways (e.g., trachea) compared to the high generations that are deep into the lungs. This aligns with the clinical observation that the possibility of initial infection is highly likely around the mouth, nose and throat [2]. Fig. 6(*III*) presents a simulated end-to-end concentration evolution of the virus over a number of days to illustrate the effects of the infection and replication process once the virus has been deposited into the airway. This concentration evolution is based on Eqn. 14. Our results shows that the levels of the virus concentration reaches its peak, on average, within 5-7 days after the initial infection, and reduces as the immune response aggressively impacts on the virus. This prediction is in line with clinical observations discussed in the study in [17]. Thus, Fig. 6(*III*) justifies the need for self-isolation between 7-14 days post-infection as the virus concentration would be at its highest during this period, which increases the probability of onward transmission.

**Figure 6:**
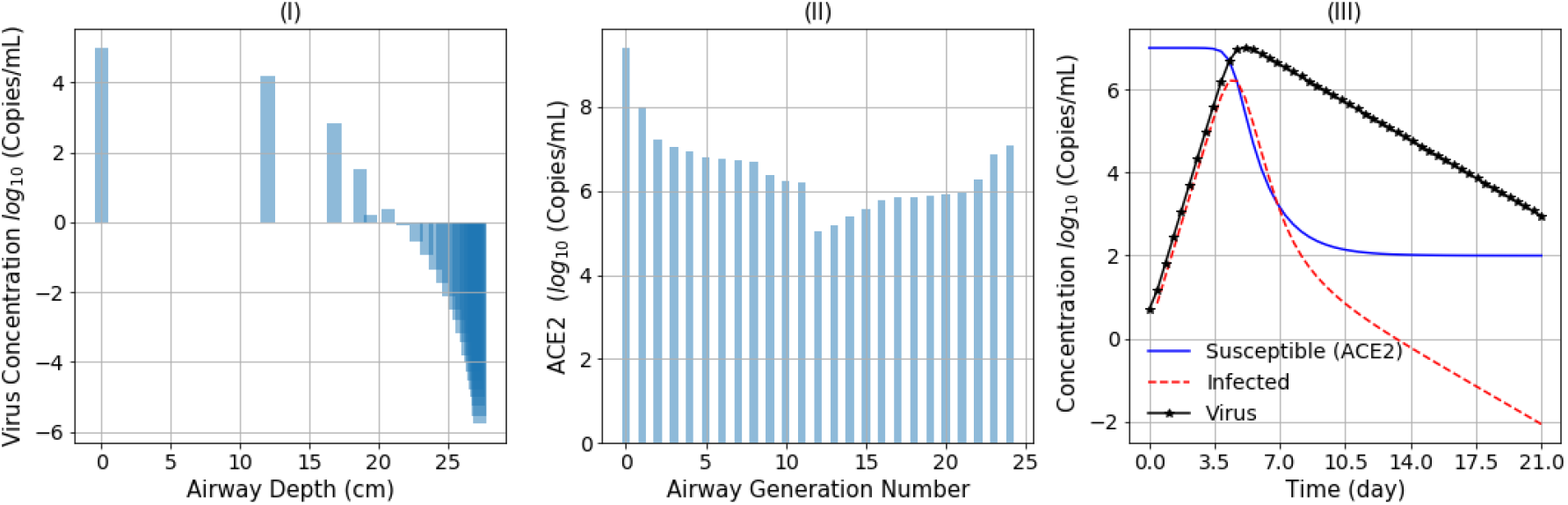
Virus deposition along the airway generation branches and infection: (*I*) its changes with respect to the initial virus load and (*II*) distribution of ACE2 expressed cells over the respiratory airways, and (*III*) simulation of viral, infection and ACE2 concentrations over time as a result of the viral infection.

The insights observed from Fig. 6(*I* − *III*) provide greater understanding of the virus deposition and infection process, and subsequently the variability in virus concentration over time. These virus concentration dynamics, however, will vary from person to person depending on a number of factors such as strength of the immune response, respiratory rate and strength, pre-existing conditions, lung airway branching structure and density of cells expressing ACE2. Therefore, to understand the impact of some of these factors, the study further explored the impact of the immune response (Fig. 7a), ACE2 expression distribution (Fig. 7b), and airflow rate (Fig. 7c) on changes to the virus load along the respiratory tract. Fig. 7a shows that the virus load decreases with increasing immune response and in particular at the lower generation levels of the respiratory track. In the case of a low immune response level, the concentration of virus still decreases moving through the lung generation levels until the highest generations where concentrations rise steeply. This is the deep region of the respiratory tracks in the alveolar region that contains high density of ACE2 expressed cells. This shows the virus particles that have not been blocked properly by the immune system, will propagate deeper into the lungs and replicate at a high rate, and addresses the case of patients with low immune system who struggle to combat the virus. Fig. 7b depicts that the greater expression of ACE2 also has a significant influence on increasing the spread of virus load over the respiratory tract, where we see that ACE2-expressed cells that are evenly spread throughout the respiratory track will result in high concentration. However, low density of ACE2-expressed cells that are in the high airway generation branches of the lung will result in a low concentration of virus, largely due to the minimum opportunities for cell entry by the virus. Fig. 7c shows that the increasing air flow rate will cause an increasing in virus concentration though, it does not exhibit a noticeable impact on the distribution of viral concentration over the respiratory tract.

**Figure 7:**
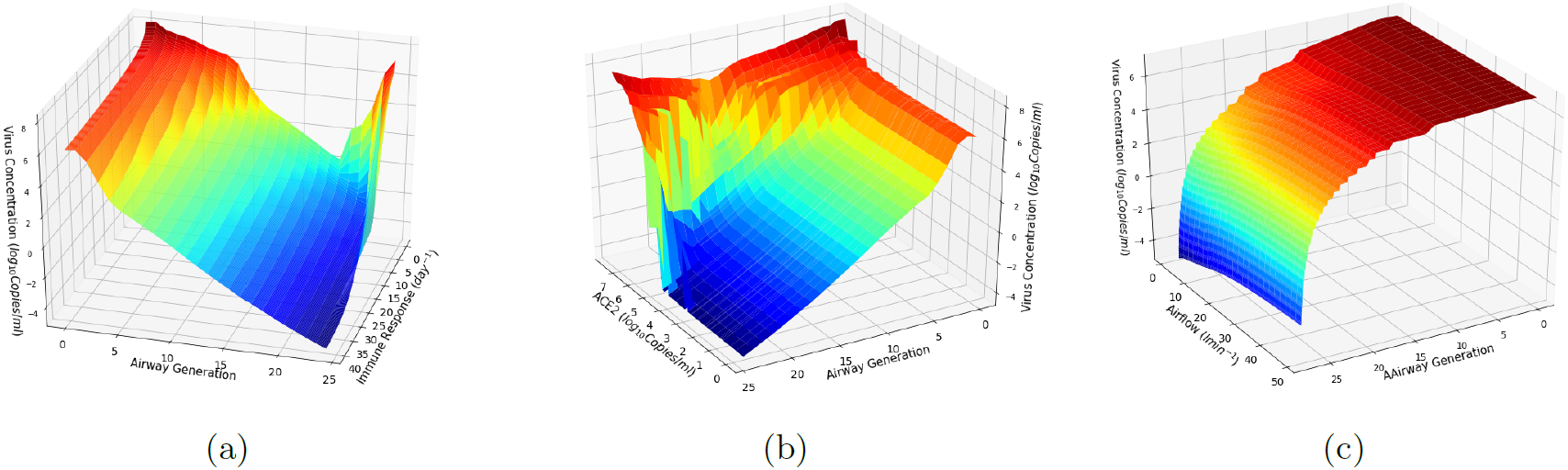
Virus load over the respiratory airways with respect to the changes in the; **(a)** immune response *d* (*Q*_0_ = 30 *lmin*^−1^), **(b)** density of ACE2 expressed cells (*Q*_0_ = 30 *lmin*^−1^ and *d* = 4.71 *day*^−1^), and **(c)** inlet airflow rate *Q*_0_ (*d* = 4.71 *day*^−1^); where the virus concentration is computed while varying the virus particle diameter *d*_*p*_ over the range 70-140 *µm* and fixed initial virus concentration *C*_0_ = 10^7^ *Copies/ml*.

### 3.3 Temporal Variability in Virus Load Over the Respiratory System

As presented in Fig. 6, the virus concentration dynamic starts as soon as the viral particles enter into the airways and start the infection process. Subsequently, the infected cells contribute to the virus concentration while the hosts innate immune system seeks to inhibit it creating a dynamically changing virus concentration environment. This process will evolve over time, with initial dose of virus as well as the effectiveness of the immune response being key influencers of the outcome. Assuming that there is constant virus reproduction and immune response rate throughout the airway generations, Fig. 8 represents the change in the virus load along the respiratory tract over time. As observed in this graph, the time to reach the peak virus load increases with the airway generations. The virus will continuously flow downwards into the lower parts of the respiratory tract, thus amplifying any replication that is taking place in the lung. Moreover, the virus load decreases after reaching the peak as susceptible cells reduces and the immune response increases. In addition, Fig. 5 in the SI 4.4 shows the variability in the virus load per airway generation in the respiratory tract.

**Figure 8:**
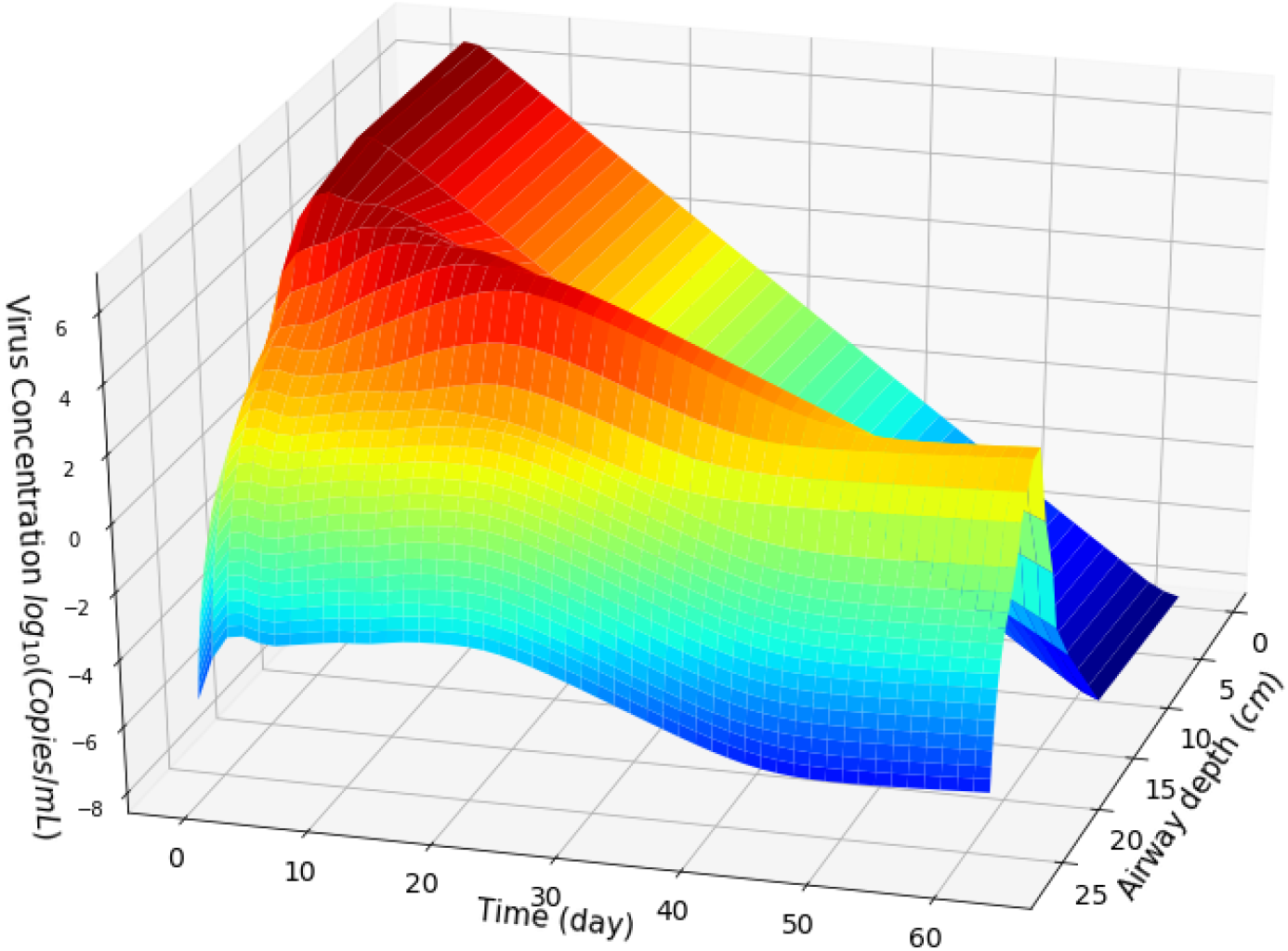
Variability in the virus load in the respiratory airway generations over time, given that the initial airflow rate is *Q*_0_ = 30*lmin*^−1^, virus particle diameter *d*_*p*_ = 70*nm*, and virus exposure quantity in contact is *C*_0_ = 10^7^*Copies/ml*.

### 3.4 Impact of Airflow Rate, Immune Response and Exposure Level on the Temporal Virus Concentration Dynamics

Eqn. 2 and 14 accounts for the impact of a number of parameters that could make a significant impact on the temporal virus concentration dynamics in the respiratory track. We have mentioned earlier that the spatial impact of these parameters on the spread of virus over the various levels of generation branches will vary from person to person, depending on different personal parameters.

To explore the impact of these parameters on the changing temporal viral dynamics, Fig. 9 presents the variability in the temporal virus behavior with respect to the change in three selected parameters, which are the inlet flow rate (*Q*_0_), immune response rate (*d*) and virus exposure level (*C*_0_). Fig. 9a(*I* − *III*) depict the change in the virus load over time for increasing three immune response *d* and flow rate *Q* values, while the virus exposure level *C*_0_ is fixed to 10^8^ *Copies/ml*. Fig. 9b(*I* − *III*) present the temporal virus behavior with respect to the change in the virus exposure level *C*_0_ and inlet airflow rate *Q*_0_ for fixed immune response of 4*day*^−1^, and Fig. 9c(*I* − *III*) depict the impact of the virus exposure level and immune response on the dynamic virus concentration levels along the respiratory tract when the airflow rate is 30*lmin*^−1^ (the exposure level *C*_0_ is presented in *log*_10_ scale). Comparing Fig. 9a(*I*) with Fig. 9a(*III*), it can be observed that the spread of virus along the respiratory tract is higher for low immune response, whereas Fig. 9b(*I* − *III*) depict that the chance of the virus propagating down the respiratory tract is high with high virus exposure level. However, in both cases the breathing rate *Q*_0_ does not seem to have a significant impact on the change in virus load. For instance, the virus load is nearly higher than 10^5^ *Copies/ml* only in the first few airway generations (∼0-3 generations) when the initial exposure is 10^1^0*Copies/ml* and the inlet airflow rate is 15 *lmin*^−1^ in Fig. 9b(*I*). However, when the initial exposure is 10^4^*Copies/ml* and the inlet airflow rate is 60 *lmin*^−1^ as presented in Fig. 9b(*III*), the virus load is nearly around 10^5^ *Copies/ml* in most of the airway generations. Considering the impact of the immune response *d* along with the exposure level *C*_0_ on the temporal behavior of the virus in the airways, Fig. 9c(*I* − *III*) shows that although the virus load that propagates down the airways decreases with higher immune response, the exposure level has a considerable impact on increasing the level of the virus load that propagates down the respiratory tract. Therefore, based on the temporal virus propagation presented in Fig. 9 and despite an increase in the inlet airflow rate *Q*_0_, the exposure level *C*_0_ and higher immune response *d* can have a high influence on the spread of the virus deep down the respiratory airways. In order to get a more in depth understanding about the impact of these parameters on the virus behavior in the respiratory tract, please see Fig. 6 in the SI 4.4.

**Figure 9:**
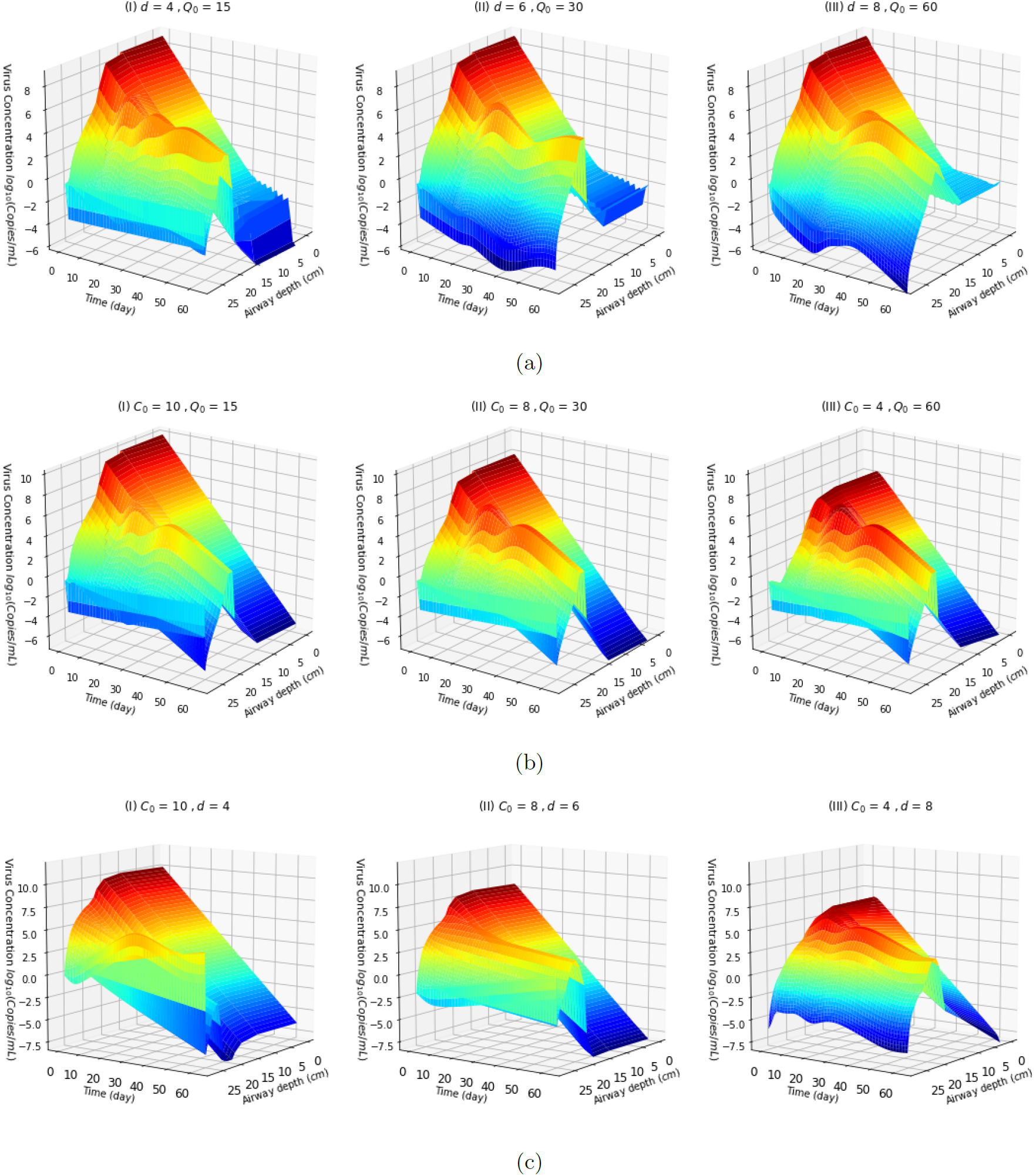
Virus load over the respiratory airways with respect to the change in; **(a)** immune response *d* and airflow rate *Q*_0_ for fixed exposure level *C*_0_ = 10^8^ **(b)** exposure level *C*_0_ and airflow rate *Q*_0_ for fixed immune response *d* = 4*day*^−1^, and **(c)** exposure level *C*_0_ and immune response *d* for fixed airflow rate *Q*_0_ = 30*lmin*^−1^ ; where virus particle diameter *d*_*p*_ varies over the range of 70-140 *µm*.

### 3.5 Impact of Effective Dose

Effective exposure level that is required for initiating the infection is still a debatable factor due to the lack of clinical evidence. The computational model investigated in this study based on the model in Eqn. 3 suggests that the virus load along an airway increases with higher exposure levels. Nonetheless, the chance of initiating an infection with a small dose of virus will likely be small as the immune system can effectively fight against the virus based on Eqns. 14-16. However, in the event of high exposure to the virus, the immune system may not be strong enough to fight against the virus replication, and this increases the probability of the infection reaching the lungs.

Fig. 10(*I* − *III*) exhibit the variability in virus concentration over airway generations for three different immune response *d* values while the exposure level *C*_0_ increases from 1 to 10000 *Copies/ml* given that the respiratory rate is *Q*_0_ = 30 *lmin*^−1^ (*d* = 0.0 means *d* = 10^−5^). The virus load propagates to the lower airway generations increases with higher exposure level, but it decreases when the immune response gets stronger (i.e., *d* increases from 10^−5^ to 10). For instance, when the exposure level is 1 *Copies/ml*, Fig. 10(*I*) shows that the virus load is *>* 1 *Copies/ml* above the 15th generation, whereas that is around 0-5 *Copies/ml* when the exposure level in 10^4^ in Fig. 10(*III*). This situation may be in line with a healthy patient having mild symptoms only. Nonetheless, when the exposure level is significantly higher than the strength of the immune response (e.g., *C*_0_ = 10^5^ *>>* 10^1^ = *d*) as depicted in Fig. 10(*III*), a larger quantity of viral particles propagate along the respiratory tract regardless of the impact of the immune response. This in turn, will increase the risk of lung infection and development of severe symptoms such as pneumonia. Therefore, when the exposure level is approximately greater than 100 *Copies/ml*, there is a greater chance of developing severe symptoms and that development may further accelerate with a weak immune system.

**Figure 10:**
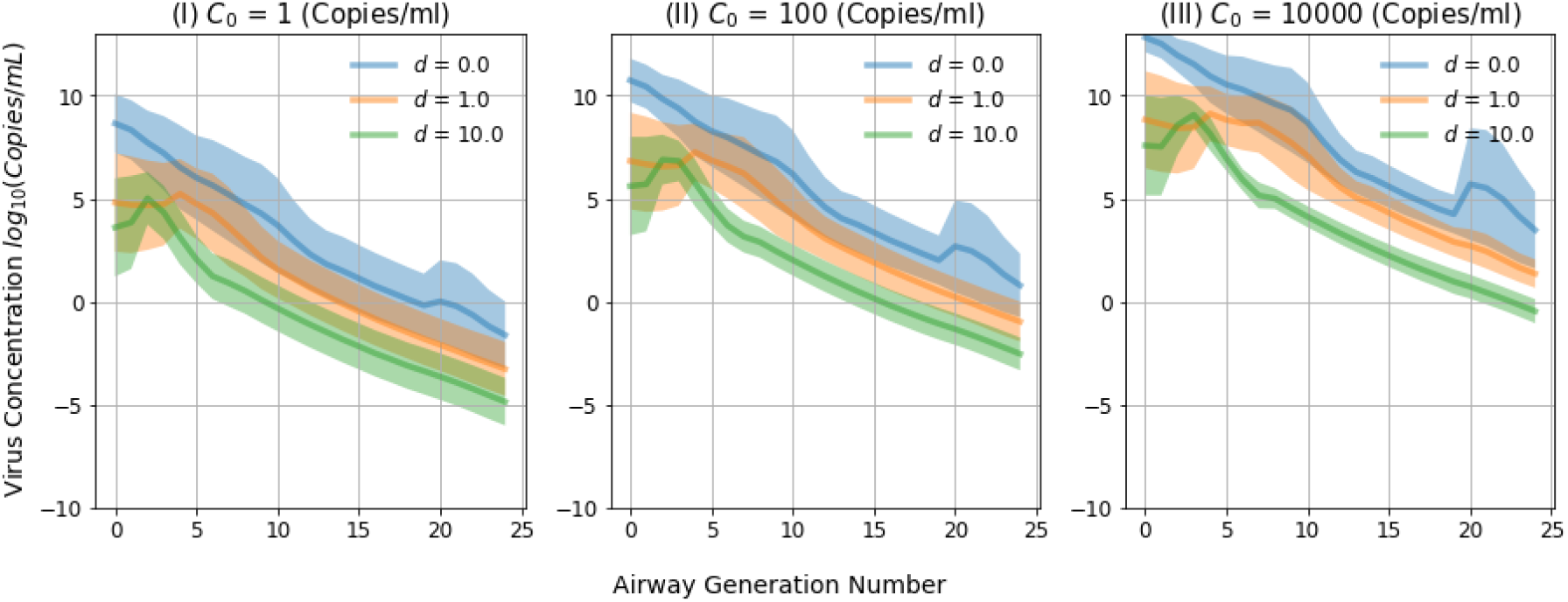
Variability in the virus load in the respiratory airway generations with respect to three different exposure levels (*C*_0_) and immune response *d* values over a period of 30 days given that the initial airflow rate *Q*_0_ = 30*lmin*^−1^, virus particle diameter *d*_*p*_ = 70*nm* (here *d* = 0.0 means *d* = 10^−5^).

## 4 Discussion

The computational model presented in this study provides deep insights into the evolution of any infectious respiratory virus along the respiratory tract given information such as respiratory rate, airway structure and initial virus infection behavior. This model is, however, not limited to respiratory infections only but can also be used in several applications to understand the different issues that may emerge. Therefore, we discuss the use of this model for understanding respiratory infection SARS-CoV-2 as well as some other applications.

In response to the global COVID-19 pandemic, there has been considerable research undertaken to understand the virus, screen current and novel compounds to identify drugs that might be effective against SARS-CoV-2 and to develop vaccines (e.g., the study [18] identified multiple drugs that were effective against SARS-CoV-2 and are now entering in vivo trials). This study reports the development of a model that maps the evolution of virus concentration dynamics throughout the respiratory tract with respect to a range of parameters that affect the propagation of the virus.

### 4.1 Tracking Temporal Propagation of Viral Particles

The virus load and the duration of virus replication are two important factors in terms of virus transmission and infection,. The virus evolution with respect to time and generation levels within the respiratory track depicted in Fig. 8, can be used to gather valuable insights to determine the virus shedding time as well as the location for drug targets to minimize replication. A potential application of this computer model, is drug targeting for diabetic patients who are known to have abnormal immune response due to the damaged vascular endothelial cells that results in lower immune response. The recent study [19], highlighted that there is a high risk of aggravation from the viral infection in the lower respiratory tract for diabetic patients. In determining treatments for such cases, knowledge about the virus propagation along the tract is the most critical information that is required in understanding the evolution of the viral infection. Deriving such insights through clinical studies is not always possible. For instance, the natural kinetics of SARS-Cov-2 in the human lungs is not fully understood [20]. Generating such valuable insights to decide effective treatments by using the computational model presented in this study can provide advantages that are difficult to be acquired through clinical practices. For instance, given certain data on the virus load in the nasal passage for a certain period of time, the model is capable of generating a complete image of the predicted virus load along the respiratory and even along each airway generation at any stage of the infection. The information about the virus evolution may, therefore, be incorporated to effectively decide the optimal drug potency required to suppress viral replication along specific branches on the airway.

### 4.2 Virus replication and propagation

Based on the mathematical models of airflow dynamics using the Navier Stokes equations, the probability of the viral particle propagation and deposition deep into the lungs is high due to their micrometer scale diameter. However, our study has found a number of factors that affects the propagation dynamics, which fluctuates as it flows through the branch generations of the airways. It is well known that the initial viral infection occurs in the upper respiratory system and then spreads further down into the lung [18]. The main reason behind this is because the virus binds to cells expressing ACE2. Fig. 4b shows the deposition rate is much higher in the upper airways under the impaction and sedimentation process. This is due mainly to the propagation of virus under the advection mechanism, which encounters turbulence in the airflow leading to increased collision of viral particles on the airways walls. Fig. 6(*I*) also shows that the majority of viral particles deposit over the upper respiratory tract, and this leads to a faster replication process before traveling down deep into the lungs as shown in Fig. 8 (this also shows that the majority of the viral transmission due to exhaling occurs due to the high replication rate that can occur in the upper respiratory tract close to the nasal passage). There is also resistance against the flow of the viral particles due to the cilia, hair and mucus lining along the upper respiratory tract; in our analytical model, the resistance due to the cilia, hair and mucus lining was accounted as the drag coefficient and slip correction factor in the computation of settling velocity *u*_*g*_ under Eqn. 34 in SI 4.4. Fig. 4a shows the decrease in the viral particle velocity due to these obstructions and this also results in faster deposition rate that is dominated by the diffusion process deep down the lungs. As with most viral proliferation process, Fig. 6(*III*) shows that the replication process can lead to 100-1000 fold increase in virus particle number during the initial 24 hours of an infection. This replication rate is compromised as the immune system starts to attack the virus based on the innate immune response first line of defence [21]. In our study, the virus replication was derived by numerically solving the Eqn. 14 with clinical viral infection data of the inlet virus concentration, immune response rate, concentration of ACE2-expressed cells as well as the viral dying rate in [2, 18]. Due to the minimal amount of information available on the replication rate of ACE2 expressed cells, our analytical model assumes that the replication rate occurs at the same rate though, certain studies have shown that the replication rate varies between different cells [22, 23]. As more data becomes available the model can be rapidly adapted to account for this.

It is clear from this model that ACE2 expression is a key factor that determines virus infection, replication and distribution in the respiratory system. ACE2 expression distribution dominates the viral infection and replication, among all the other parameters. It is known that a number of factors such as smoking [24], age, gender and race [25] can lead to varying densities of ACE2-expressing cells, and these differences will have a key impact on clinical outcome as well as how easily a person will transmit the virus to others. Our analytical model considers variability in the density of ACE2-expressed cells along the branches of the airway tract, where there is greater ACE2-expressed cell density in the nose and mouth (airway generation 1-2) as well as in the alveoli region (airway generation 21-24). This complies with a recent study [9] that suggested the oral mucosa has good expression of ACE2, while the study [26] highlighted that the ACE2-expression seems predominantly greater in the AT2 (type II alveolar) cells that express high levels of ACE2 in the lung, while the upper airways have a broad distribution of ACE2. This is why the virus load is considerably higher in the top and lower sections of the respiratory track compared to branches in the middle airway generations, and this is shown in in Fig. 7a for the lower immune response values. In addition, Fig. 7b also highlights the greater possibility of viral infection being distributed over the upper airways and also alveoli region due to the higher density of ACE2 cells. Since our model is based on a spatial branching structure of the airways, as more details of ACE2-expressed cells is available for each of the branch links, this information can be inserted into our analytical model to gain more detailed insights into the proliferation process along the airway network.

### 4.3 Immune Response to Viral Proliferation

It is already known that the most vulnerable groups to the SARS-CoV-2 are the elder people and people with chronic diseases [27]. This is likely to be because because of their weakened immune system that is not as effective at fighting against invasive pathogens such as SARS-CoV-2 compared to healthy individuals. This can be attributed to a general reduction in efficiency of the immune system as well as a number of lifestyle habits such as smoking, imbalanced nutrient intake along with environmental factors such as air pollution that can affect the immune system, making individuals more vulnerable to SARS-CoV-2. Our study provides insights into the impact of immune response and its role in the viral proliferation down the branches of the respiratory tracts. Fig. 9a(*I* − *III*) and Fig. 9c(*I* − *III*) shows that stronger immune response can effectively control the spread of the virus down the respiratory tract even when a patient is exposed to a large quantity of viral particles. Therefore, the computational model can provide a prediction tool for people who are at high risk based on the effectiveness of their immune system. This model is of course focusing on the initial infection and does not take into account secondary cytokine storm type responses that follow an extended SARS-CoV-2 infection. However, it does suggest that a stronger initial immune response is particularly effective at limiting advancement of virus to the lower parts of lung where serious damage can occur.

### 4.4 Exposure Level and Effective Dose

It is believed that a healthy person can take in the virus through either direct or indirect exposure. The direct exposure occurs by being in close vicinity to the expulsion of the droplets from an infected person (i.e., coughing, sneezing, singing, talking and heavy breathing), while the indirect exposure is when the hands touch contaminated surface of droplets and this comes in contact with either the mouth or nose. Therefore, theoretically, the direct exposure has a higher risk of receiving a large dose of virus. However, it is still unclear what impact the concentration of the initial exposure dose has on clinical outcome. It is important to also note that symptoms do not necessarily correlate with virus load by nasal swab. In fact asymptomatic people have similar RT-PCR values to symptomatic people, indicating that asymptomatics can probably transmit virus as efficiently as symptomatics. The difference however, may the localization of the virus during the initial days of infection. If the initial dose is high, this may overwhelm the front line defence and move rapidly to the lung where serious disease is caused.

Hence, this study aims to also understand the effect of the exposure level on the virus load and distribution, and how this is impacted by other parameters such as the immune response and breathing rate. Fig. 9 shows that the virus load over the respiratory tract does not only depend on the initial exposure level, but also the counter effect from the immune response. However, the respiratory rate does not have a considerable influence on the spread of the virus load. For instance, Fig. 9a(*I* − *III*) shows that greater immune response can control the virus load. Moreover, Fig. 10(*I* − *III*) also depicts that the weak immune response and high virus exposure level can make a significant influence on the severity of infection. When the exposure level is significantly higher (at least greater than 100 *Copies/ml*) compared to the strength of the immune response, the chance of reaching a considerable quantity of virus in the lungs is very high, regardless of the respiratory rate.

In terms of real-world applications of this model, there are many possibilities. For example, the computational model could, therefore, serve as a tool to understand how nanoparticles that are proposed for binding on the virus, can propagate down the respiratory track and at the same time determine the optimal quantity of the nano particles to be inhaled into the patient. In particular, the computational model can be used to predict the required treatment for patients who are in the high risk category (e.g., smokers). Moreover, the model also helps us to predict the infective capabilities of future viruses that jump the species barrier, thus improving out ability to be ready for another potential pandemic.

## Acknowledgement

This research was supported by a research grant from Science Foundation Ireland and the Department of Agriculture, Food and Marine on behalf of the Government of Ireland under the Grant 16/RC/3835 (VistaMilk).

## Supplementary Information

### Virus Concentration Model Derivation 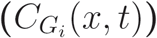

To derive a model for characterizing the virus propagation along a respiratory airway generation (*G*_*i*_), consider a control volume *V* of depth *δx* with a cross sectional area *A* as illustrated in Fig. 11 (i.e., *V* = *Aδx*). As Fig. 11 illustrates, when virus particles enter to an airway (shown as gray circles), they deposit on the airways walls and subsequently the virus infection takes place. As a consequence of the virus infection, more virus particles are reproduced (blue circles). The number of virus particles in the control volume (i.e., virus concentration, 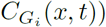 can be expressed as 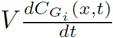. Then, the accumulation of the virus in the control volume *V* can be formulated by considering the mass balance, i.e., virus particle in-flux (*f*_*in*_) and interior flux-loss to the airway surface (*f*_*loss*_) and out-flux (*f*_*out*_) of virus particles as follows:

**Figure 11:**
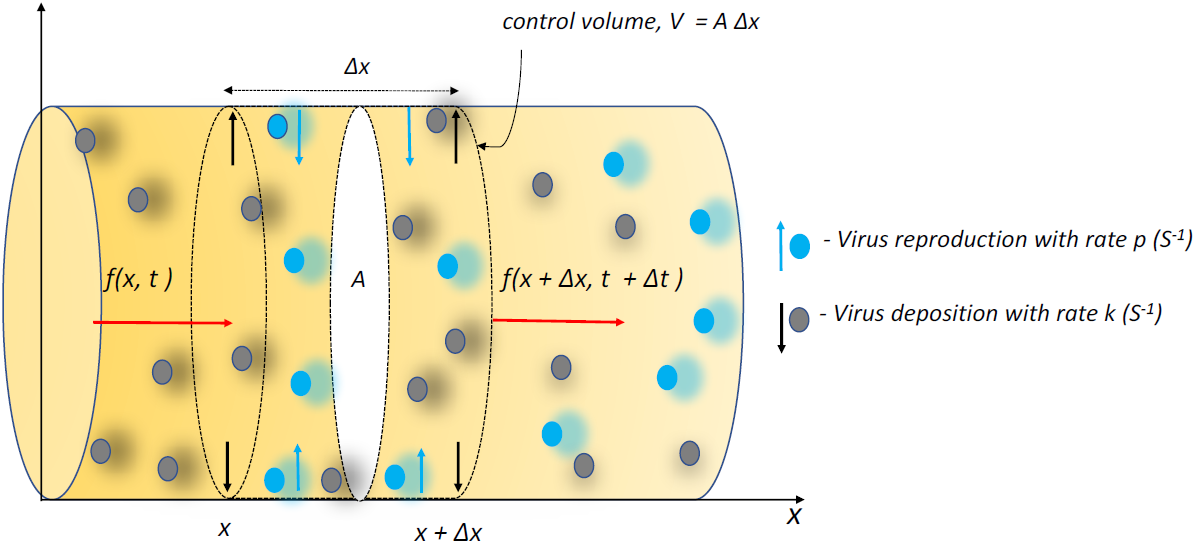
Virus particle propagation through a control volume of an airway generation with deposition rate *k* and reproduction rate *p*.

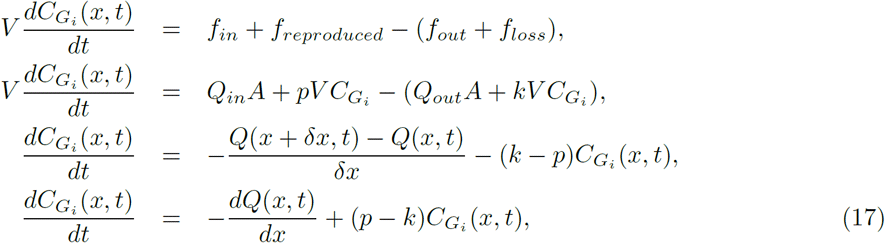

where *k*(*s*^−1^) and *p*(*s*^−1^) are respectively the virus deposition rate on the airway surface and virus reproduction rate and *Q* is the airflow rate through the control volume *Q*_*in*_ = *Q*(*x, t*) and *Q*_*out*_ = *Q*(*x* + *δt*). Since the virus particles flow under the advection and diffusion mechanisms, the airflow rate *Q* can be formulated as the total airflow due to advection and diffusion, and hence,

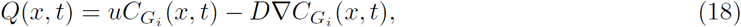

where *u* is the air flow velocity and *D* is the diffusion coefficient. Hence, the rate of change of air flow rate along the *x*-axis (in practice, *x*−axis implies the direction to which air flows), 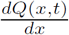, can then be written as

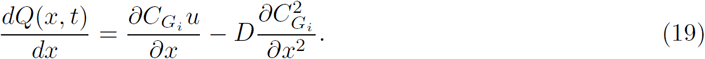

When the airflow achieves its settling velocity (i.e., when the airflow flow reaches its steady state), 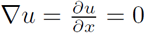 and *D* is a constant. So, Eqn. 19 can be simplified into the following expression:

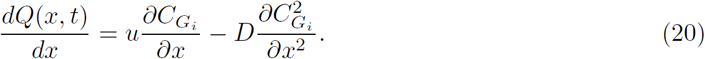

Based on the mass balance of the fluid flow without any reproduction and absorption of virus in Eqn. 17, the following relationship in Eqn. 21 holds regardless if the flow is due to advection or diffusion, and is represented as [6].

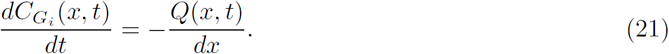

By plugging Eqns. 20 and 21 in Eqn. 17), the change in virus concentration along an airway can be represented in Eqn. 22. This is normally called the governing equation that represents the dynamics in virus concentration along the respiratory tract, and is represented as

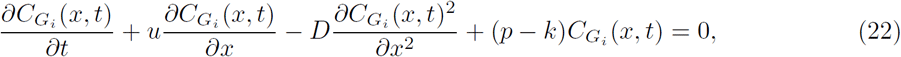

for *i* = 0, 1, *…*, 23. Suppose the initial inlet virus concentration is 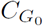, i.e., 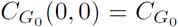, then the solution for Eqn. 22 can be derived by applying the following transformation:

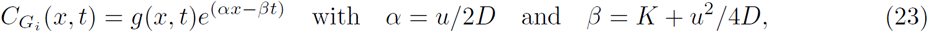

where *K* = *k* − *p*. Then the partial derivatives of Eqn. 23 with respect to time *t* and *x* are

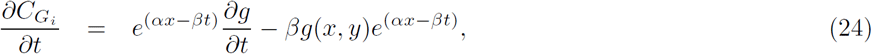

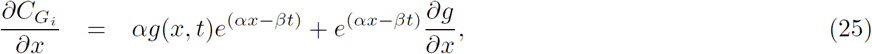

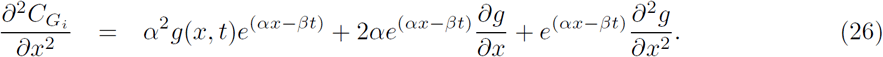

By plugging these derivatives in Eqn. 22, it can be converted in the form of the standard steady state diffusion equation in terms of *g*(*x, t*) as given in Eqn. 27:

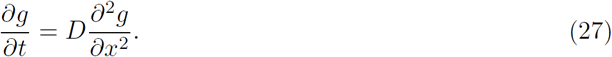

The general analytical solution of Eqn. 27, the virus concentration, 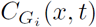, at a point *x* and time *t* can be written as

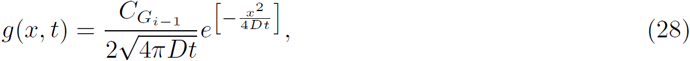

where 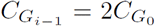 only at *t* = 0. This is because the 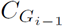 divides equally into two when virus enters into the two branching airways of the *G*_*i*_ generation.

The solution of Eqn. 22 can then be expressed as

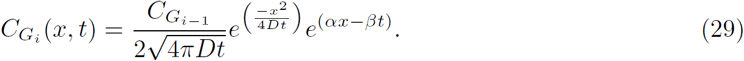

By substituting the values of *α* and *β* given in Eqn. 23 in Eqn. 29 and then re-arranging the terms in the exponential function, the final solution of Eqn. 22 can be expressed as follows

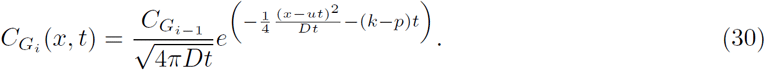

This expression presents the change in the virus concentration along an airway over time along with the effects of virus reproduction and deposition.

At the steady state, 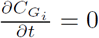 and *u* is also a constant. Then, the Eqn. 22 can be simplified as

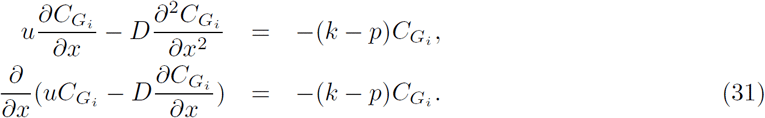

Note that according to Eqn. 18, 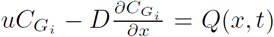. The analytical solution of Eqn. 31 can be derived by transforming it into the following quadratic equation:

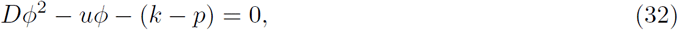

where 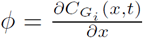. The solution of Eqn. 32 can be written as 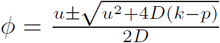. Since the concentration cannot be negative or an imaginary number, the condition *u*^2^ + 4*D*(*k* − *p*) *>* 0 should hold, that is 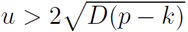. Hence, the final solution of Eqn. 32 is

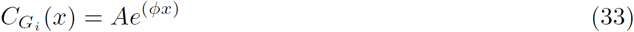

where *A* is an integration coefficient that has to be determined. To compute the value of *A* in the steady state considering Eqn. 31, this can be represented as

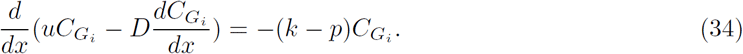

It is given in Eqn. 19 that 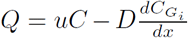. At *x* = 0, from 33, 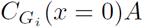 and 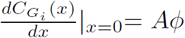 and 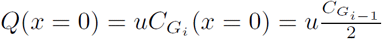. Hence, according to 34,

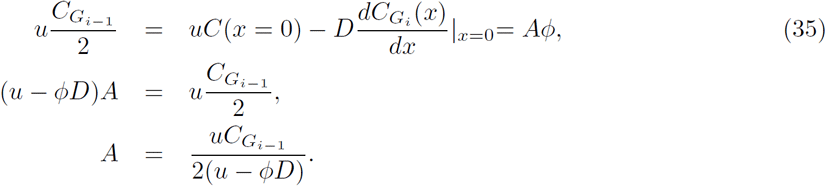

Now, *A* can have two values *A*_1_ and *A*_2_ as follows

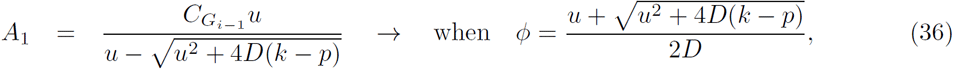

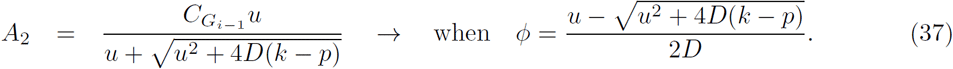

Considering Eqn. 36, *A*_1_ *≥* 1 as 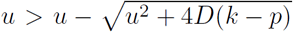, and this turns out that according to Eqn. 33, 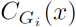 is increasing with *x*. On the other hand, according to Eqn. 37, *A*_2_ *≤* 1 as 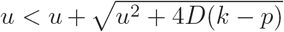, hence *C*(*x*) decreases with *x*. In practice, the virus concentration should decrease when the virus travels along the airway tract. Thus, *A*_2_ is the most suitable value for *A*.

Finally, by substituting the *A* and *φ* values derived from Eqn. 35 and Eqn. 32 in eqn. 33, the steady state virus concentration along the airway tract, 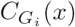, can be written as

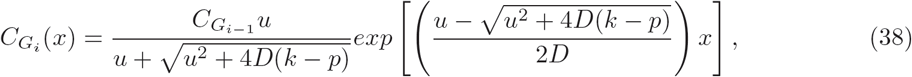

for 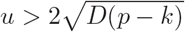 and *i* = 0, 1, *…*, 23.

### Impedance (*Z*) airflow rate (*Q*) and airflow velocity (*u*)

Based on the conservation of fluid flow, the following conditions hold at a bifurcation junction as represented in Fig. 12.

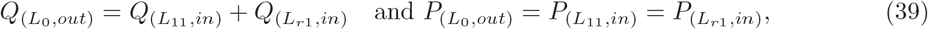

where *Q*-flow rate, *P* - pressure, *L*_*l*1_ and *L*_*r*1_ are respectively the length of the left (*l*) and right (*r*) branches of the parent airway of length *L*_0_ (or in generation 1, *G*_1_). The subscript *in* and *out* stands for the inlet and outlet of the airway, respectively.

**Figure 12:**
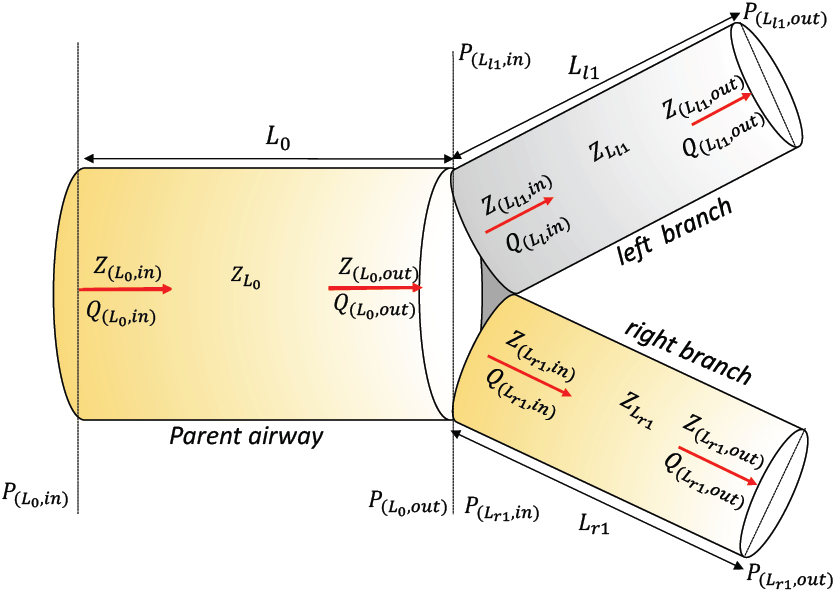
Bifurcation of the first airway generation (*G*_0_), i.e., trachea into bronchus.

Based on the circuit theory (i.e., Ohm’s law; *V* = *IR*, where *V* -voltage, *R*-resistance and *I*-current), the impedance (*Z*), resistance against the airflow, is then computed as *Z* = *P/Q* [5]. Thus, the following relationship can be derived to compute the impedance at the junction (i.e., outlet of the pipe) as

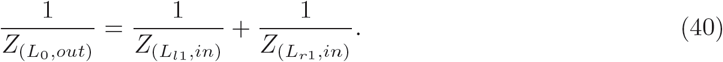

Thereby, the impedance at the inlet of the parent airway is computed as

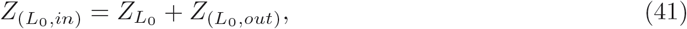

where 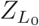 is the impedance of the parent airway (i.e., *G*_0_) and computed based on the following procedure.

Following the definition of impedance *Z*, i.e., *Z* = *P/Q*, and the concepts in fluid dynamics, the impedance of the parent airway 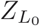 is expressed as

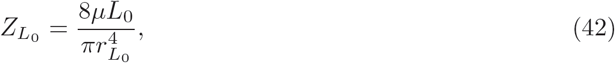

where *µ*-airflow viscosity and 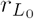 is the airway diameter. By plugging Eqn. 42 in Eqn. 41, the impedance at the inlet of the parent airway can be calculated as

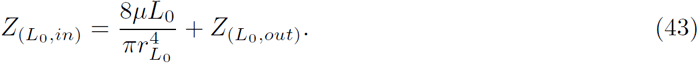

Now, at the bifurcation junction, the inlet flow rates into the left and right branches of the parent airway as well as its outlet pressure are then computed as

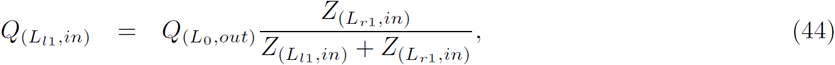

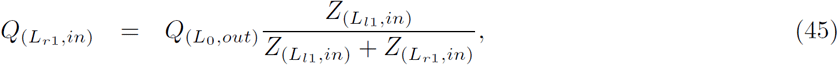

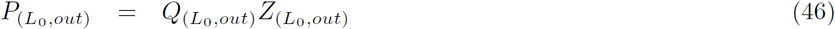

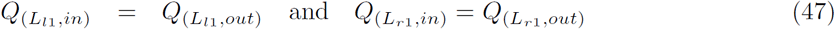

As Fig. 12 represents the bifurcation (branching) of the respiratory tract as airway generations, a backward process is followed to compute the airflow rates, and then the velocity profiles over the respiratory tract.

Having computed the impedance of each airway generation, the airflow rate *Q* is then computed by using Eqns. 44-47. Finally, the *Q* = *uA* relationship is used to compute the airflow velocity profile of each airway generation.

### Settling Velocity (*u*_*g*_)

The maximum velocity that a virus particle can achieve is known as its settling velocity; velocity of a particle when the total external forces acting on it becomes zero. Fig. 3, for instance, depicts a virus particle of mass *m*, diameter *d*_*p*_ and density *ρ*_*p*_ that enters to an an airflow of density *ρ*_*f*_ at *t* = 0 with velocity *u*_0_ and then propagates along an airway with velocity *u*_*f*_ under three external forces given in Eqns. 48-49. We assume the virus particle achieves its settling velocity *u*_*g*_ at *t* = *T* after traveling *L*_*T*_ distance. When the virus particle moves through a projected area *A*_*p*_ at an arbitrary time *t*, its relative velocity (velocity with respect to the airflow) can be expressed as *u*_*p,t*_ = *u*_*f*_ − *u*_*p*_, and it also gains an acceleration due to the imbalance of the external forces that act on it, and this includes

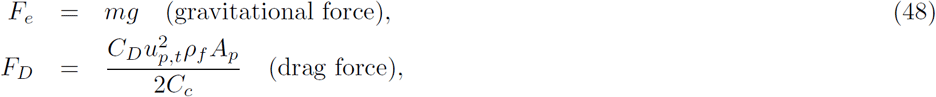

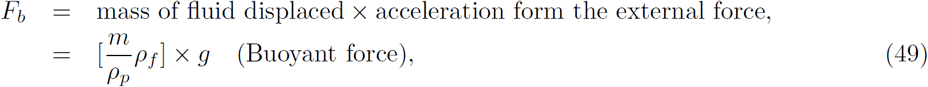

where *C*_*D*_ is the drag coefficient and *C*_*c*_ is the slipping coefficient. The Newton’s second law (i.e., 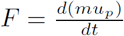 can then be used to express relationship of these three forces as

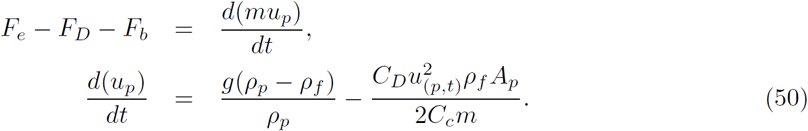

According to the definition of the Reynolds’s number, Reynolds’s number of the virus particle is 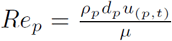 and hence the relative velocity can be expressed as 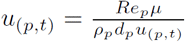. Moreover, assuming that the spherical shaped virus particles, the weight *m* and cross sectional area *A*_*p*_ of the virus particle can be expressed as 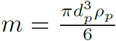 and 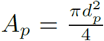, respectively. By plugging these terms in Eqn. 50, then Eqn. 51 expresses the change in particle velocity *u*_*p*_ as

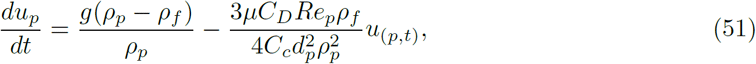

where *g* is the acceleration gravity and

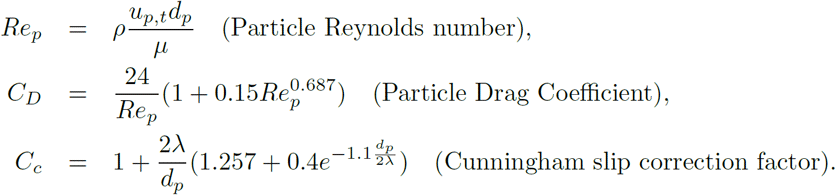

The virus particle achieves the settling velocity, *u*_*g*_, when 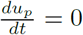 (i.e., Eqn. 51 = 0), hence the settling velocity can be expressed as

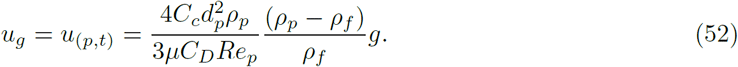

## Additional Results

Some additional results is presented here, aiming to give more in depth understanding about the viral dynamics along the respiratory tract.

### Impact of Breathing and Virus size on Virus Deposition

In addition to the impact of the airflow rate (or respiratory rate) *Q*_0_ on the flow velocity *u* and deposition rate *k* presented in Fig. 5a(*I* − *III*) and 5b(*I* − *III*) in the main document, particle size (this study considers the diameter *d*_*p*_) can also impact on the particle deposition as larger particles are mostly trapped in the upper airways, while smaller (e.g., micrometer scale) particles propagate deep down the lungs. When the airflow rate *Q*_0_ is fixed to 30*lmin*^−1^, while the impact of changes in the particle diameter on the airflow velocity is presented in Fig. 14a(*I* − *IV*), Fig. 14b(*I* − *IV*) shows the influence of the particle size on the virus deposition along the respiratory tract. The virus deposition rate increases with increasing particle size and a higher deposition can be observed over the upper airway generations (∼*G*_0_ − *G*_15_). However, the increase in virus deposition rate is significantly less in the lower airways compared to the upper airway generations.

**Figure 13:**
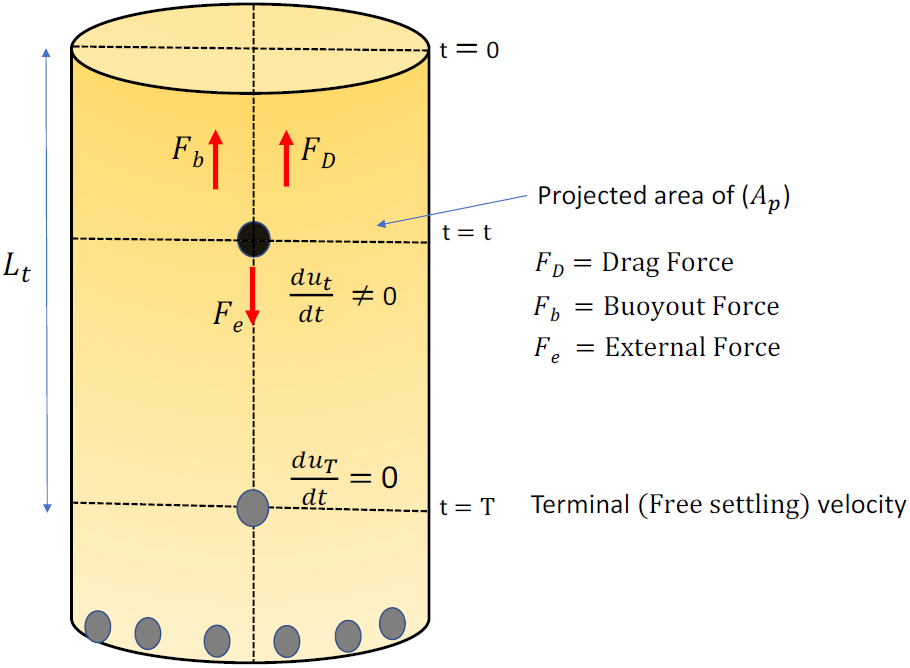
Particle propagation under gravity and settling on the airway walls.

**Figure 14:**
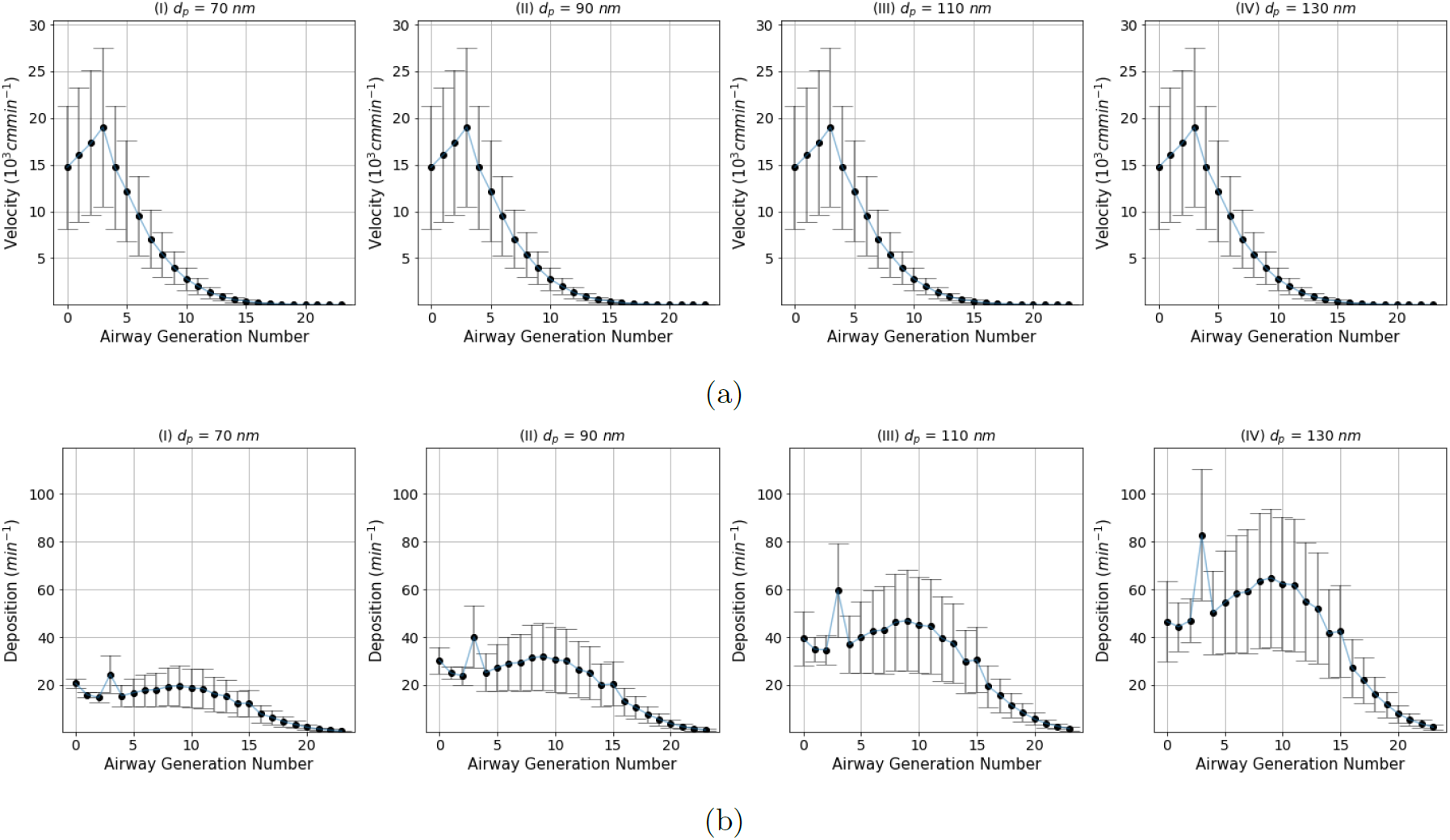
Variability in virus deposition along the respiratory tract with respect to **(a)** changing breathing rate *Q*_0_ with virus particle diameter 70*µm*) and **(b)** virus diameter *d*_*p*_ when the breathing rate *Q*_0_ = 30*lmin*^−1^.

### Variability in Virus Load Over Airway Generations

Insights about the variability in the virus load over the respiratory tract is important particularly in early diagnosis and making timely treatments to control the virus spread and development into severe illness. This is critically important in treating patients with chronic diseases like diabetes. In this regard, Fig.15 depicts the variability in the virus load over the respiratory tract after 35 days given that a person exposed to the virus with an initial dose 10^5^*Copies/ml* and breathing rate is 30*lmin*. Since the ACE2 expressed cell density is high around the mouth as well as the alveolar region, the variability in the virus load is high.

**Figure 15:**
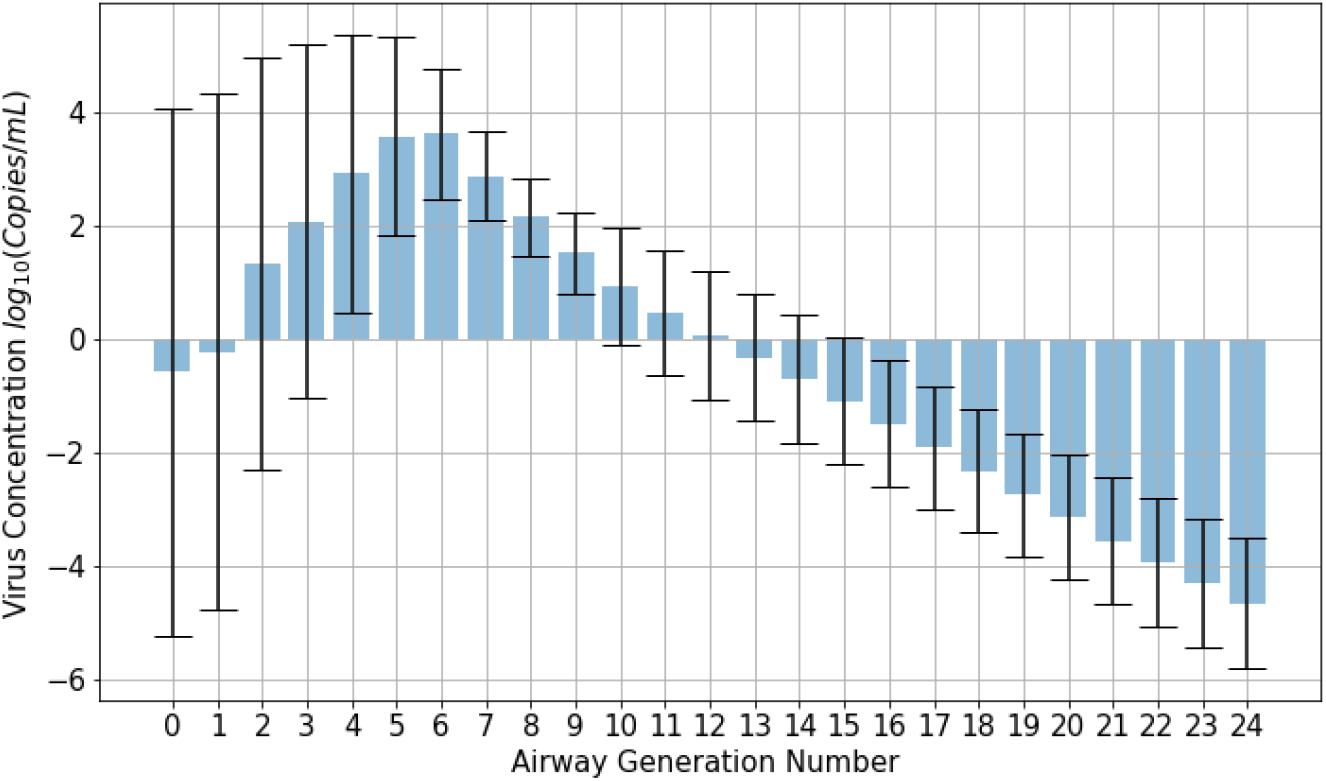
Average virus load in each of the airway generation over a period of 30 days; ACE2 concentration vary over the the upper (∼1 − 3 generations) and lower (∼*G*_21_ − *G*_23_ generations) regions as 𝒩(5 − 7, .71) and it is 𝒩 (5.83, 0.71) over the remaining airway generations, and air flow rate *Q*_0_ is 30*lmin*^−1^ and virus diameter *d*_*p*_ is 70*nm*.

### Impact of Respiratory Rate, Exposure Level and Immune Response on the Change in Virus Load

In addition to the graphs given in Fig. 9 in the main document, to understand further the temporal viral dynamics along the respiratory tract in response to the change in three selected parameters, respiratory rate (*Q*_0_), immune response (*d*) and exposure level (*C*_0_), Fig. 16 shows the temporal virus dynamics for different values of *C*_0_, *Q*_0_ and *d*. The impact of the third parameter is averaged when representing the impact of the remaining parameters on the changes in the virus load down the respiratory tract. That is, for instance, the variability in the virus load with respect to *Q*_0_ and *C*_0_ depicted in Fig. 16a(*I* − *IX*) is computed by averaging the virus load derived for the three *d* values for each *C*_0_ and *Q*_0_ combination. All in all, Fig. 16 highlights that the exposure to a higher virus load increase the chance of viral infection, although the strong immune response can suppress the spread of the virus along the respiratory tract. Similarly, Fig. 16b(*I* − *IX*) shows the impact of the exposure level *C*_0_ and the immune response *d* on changing the temporal virus dynamics along the respiratory tract and it can be observed that the virus load is considerably higher for small immune response values and higher exposure levels. Meanwhile, the impact of the respiratory rate and immune response on the variability in the virus load along the respiratory tract depicted in Fig. 16c(*I* − *IX*) highlights that when the immune response is weak, the possibility of the virus spreading towards the lungs is higher regardless of the breathing rates.

**Figure 16:**
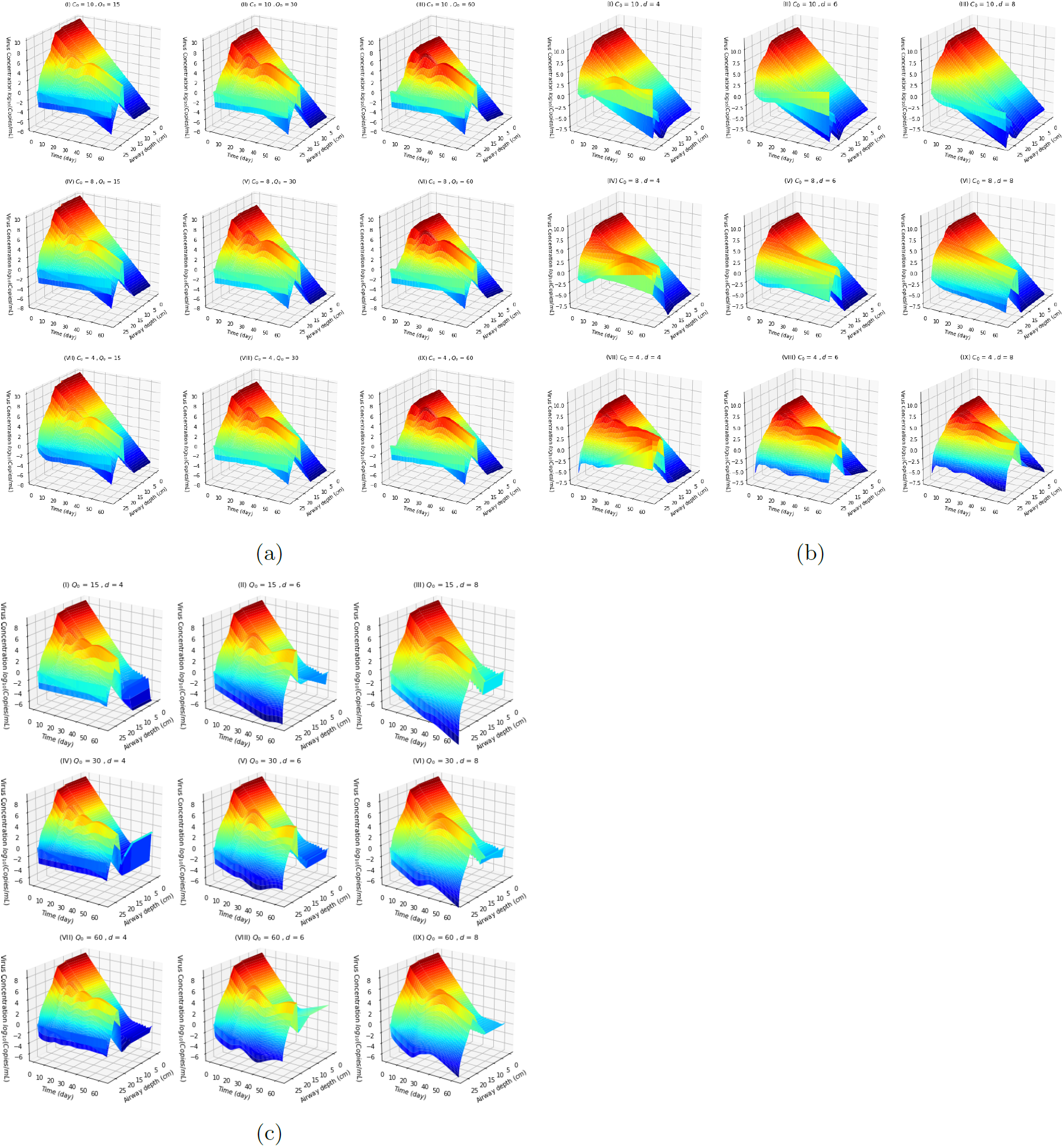
Virus load down the respiratory tract with respect to the change in; **(a)** Flow rate and exposure level (*Q*_0_ and *C*_0_) **(b)** Immune response and explore level (*d* and *C*_0_) and **(c)** Immune response and flow rate (*d* and *Q*_0_); where virus particle diameter *d*_*p*_ varies over the range 70-140 *µm*.

## Notes

### Competing Interest Statement

The authors have declared no competing interest.

